# Multimodal profiling reveals tissue-directed signatures of human immune cells altered with age

**DOI:** 10.1101/2024.01.03.573877

**Authors:** Steven B. Wells, Daniel B. Rainbow, Michal Mark, Peter A. Szabo, Can Ergen, Ana Raquel Maceiras, Daniel P. Caron, Elior Rahmani, Eli Benuck, Valeh Valiollah Pour Amiri, David Chen, Allon Wagner, Sarah K. Howlett, Lorna B. Jarvis, Karen L. Ellis, Masaru Kubota, Rei Matsumoto, Krishnaa Mahbubani, Kouresh Saeb-Parsy, Cecilia Dominguez-Conde, Laura Richardson, Chuan Xu, Shuang Li, Lira Mamanova, Liam Bolt, Alicja Wilk, Sarah A. Teichmann, Donna L. Farber, Peter A. Sims, Joanne L. Jones, Nir Yosef

**Affiliations:** Department of Systems Biology, Columbia University Irving Medical Center, New York, NY; Department of Microbiology and Immunology, Columbia University Irving Medical Center, New York, NY; Department of Clinical Neurosciences, University of Cambridge, Cambridge, UK; Department of Systems Immunology, Weizmann institute, Rehovot, Israel; Department of Electrical Engineering and Computer Science and Center for Computational Biology, University of California, Berkeley, CA; Wellcome Sanger Institute, Wellcome Genome Campus, Hinxton, Cambridge, UK; Department of Surgery, Columbia University Irving Medical Center, New York, NY; Department of Surgery, University of Cambridge, Cambridge, UK; Theory of Condensed Matter, Cavendish Laboratory, University of Cambridge, Cambridge, UK; Department of Biochemistry & Molecular Biophysics, Columbia University Irving Medical Center, New York, NY

**Keywords:** Single cell genomics, Lymphocytes, myeloid cells, macrophages, tissue residence, mucosal immunity, immune senescence, aging, tissue atlas, counterfactual analysis

## Abstract

The immune system comprises multiple cell lineages and heterogeneous subsets found in blood and tissues throughout the body. While human immune responses differ between sites and over age, the underlying sources of variation remain unclear as most studies are limited to peripheral blood. Here, we took a systems approach to comprehensively profile RNA and surface protein expression of over 1.25 million immune cells isolated from blood, lymphoid organs, and mucosal tissues of 24 organ donors aged 20-75 years. We applied a multimodal classifier to annotate the major immune cell lineages (T cells, B cells, innate lymphoid cells, and myeloid cells) and their corresponding subsets across the body, leveraging probabilistic modeling to define bases for immune variations across donors, tissue, and age. We identified dominant tissue-specific effects on immune cell composition and function across lineages for lymphoid sites, intestines, and blood-rich tissues. Age-associated effects were intrinsic to both lineage and site as manifested by macrophages in mucosal sites, B cells in lymphoid organs, and T and NK cells in blood-rich sites. Our results reveal tissue-specific signatures of immune homeostasis throughout the body and across different ages. This information provides a basis for defining the transcriptional underpinnings of immune variation and potential associations with disease-associated immune pathologies across the human lifespan.

## INTRODUCTION

The immune system comprises a complex network of specialized cells in diverse tissue sites to protect against infections and cancer, regulate inflammation, and resolve tissue damage or injury. In humans, there is tremendous heterogeneity in immune responses across tissues and over age, yet the underlying mechanisms remain difficult to ascertain.

Myeloid lineage cells, such as macrophages, monocytes, and dendritic cells, play a crucial role in coordinating innate immunity by managing inflammatory responses and antigen processing directly in tissue sites of pathogen encounter such as lungs, intestines, skin and other sites. The subsequent induction of adaptive immunity involves the activation of antigen-specific T and B lymphocytes in lymphoid organs thereby promoting B cell differentiation to antibody secreting cells, and T cell differentiation to effector cells, which migrate to infection sites for pathogen clearance. Immune memory develops from activated lymphocytes and persists as non-circulating, tissue resident memory T cells (TRM) ^1^, memory B cells in both lymphoid and mucosal sites, and circulating memory populations across sites.

However, the identity and heterogeneity of human immune cells across lineages, tissues, and age remain poorly understood, as most studies of human immunity are confined to the sampling of peripheral blood ^2^ or to multi-tissue analysis of restricted cohorts ^3^. Recent advances in single-cell technologies for transcriptome profiling have enabled comprehensive identification of individual cell types, their function, and heterogeneity in tissues throughout the body. Cellular atlases specific to organs such as lungs ^4^, liver ^5^, brain ^6^, and reproductive tract ^7^ provide comprehensive datasets for elucidating developmental pathways and serve as new baselines for understanding tissue-specific disease states.

For immune cells, investigating many tissues across diverse ages is necessary to encompass the entire breadth of the immune system and identify sources of variation ^8^. We have pioneered the study of human immune cells across diverse anatomic sites using tissues obtained from organ donors, enabling phenotypic, functional, and single cell profiling of immune cells from multiple tissues within and between individuals ^9–13^. Our studies of lymphocytes revealed that T and NK cell subset composition, tissue residence, and functional attributes are specific to the tissue ^9,12,14–16^, suggesting that tissue may play a dominant role in immune cell states across lineages and their variation with age.

In this study, we sought to define the role of tissue and age in immune cell identity and function, accounting for all major immune lineages. We used cellular indexing of transcriptomes and epitopes (CITE-seq) to simultaneously profile transcriptome and surface marker expression of myeloid and lymphoid-lineage cells in 14 tissue sites of 24 organ donors, aged 20-75 years. Through multimodal classification, probabilistic modeling, and various comparative analyses, we reveal how certain lineages and subsets exhibit tissue-specific adaptations, and how aging affects the cellular and molecular composition of immune lineages across and within tissues.

## RESULTS

### Multimodal profiling and classification of immune lineages across sites and ages

We isolated mononuclear cells from blood, lymphoid organs, lungs, airways, intestines, and other sites, using well-established protocols (see Methods)^12,13^ (**Fig. 1A, B**). We performed scRNA-seq on 24 donors using the 10x Genomics platform, of whom 22 had CITE-seq–profiling with at least 130 proteins (see Methods, **Supplementary Table 1**). The organ donors originated from New York City (US) and Cambridge, England (UK), and were free of chronic infection, cancer, and overt disease (**Supplementary Table 2**). After stringent quality control, we obtained 1.27 million immune cell events from 14 tissue sites. This included ten key sites, each with at least 75,000 cells, including blood (BLO), bone marrow (BMA) spleen (SPL), lungs (LNG) which include bronchoalveolar lavage (BAL) and lung parenchyma (PAR), lymph nodes (LN) including lung-associated (LLN), mesenteric (MLN), and iliac (ILN), and jejunum (JEJ) divided into the jejunal intraepithelial layer (JEL), and jejunal lamina propria (JLP), and these sites were subsequently analyzed as described below (**Fig. 1B**). When available, immune cells were also purified from liver (LIV), skin (SKN), colon epithelium (CEL) and lamina propria (CLP). While these data will be made available as part of our annotated dataset, they, along with polymorphonuclear cells (mast cells and neutrophils), were not included in subsequent analyses due to low cell and donor numbers **(Extended data Fig. 1A**).

**Figure 1.**
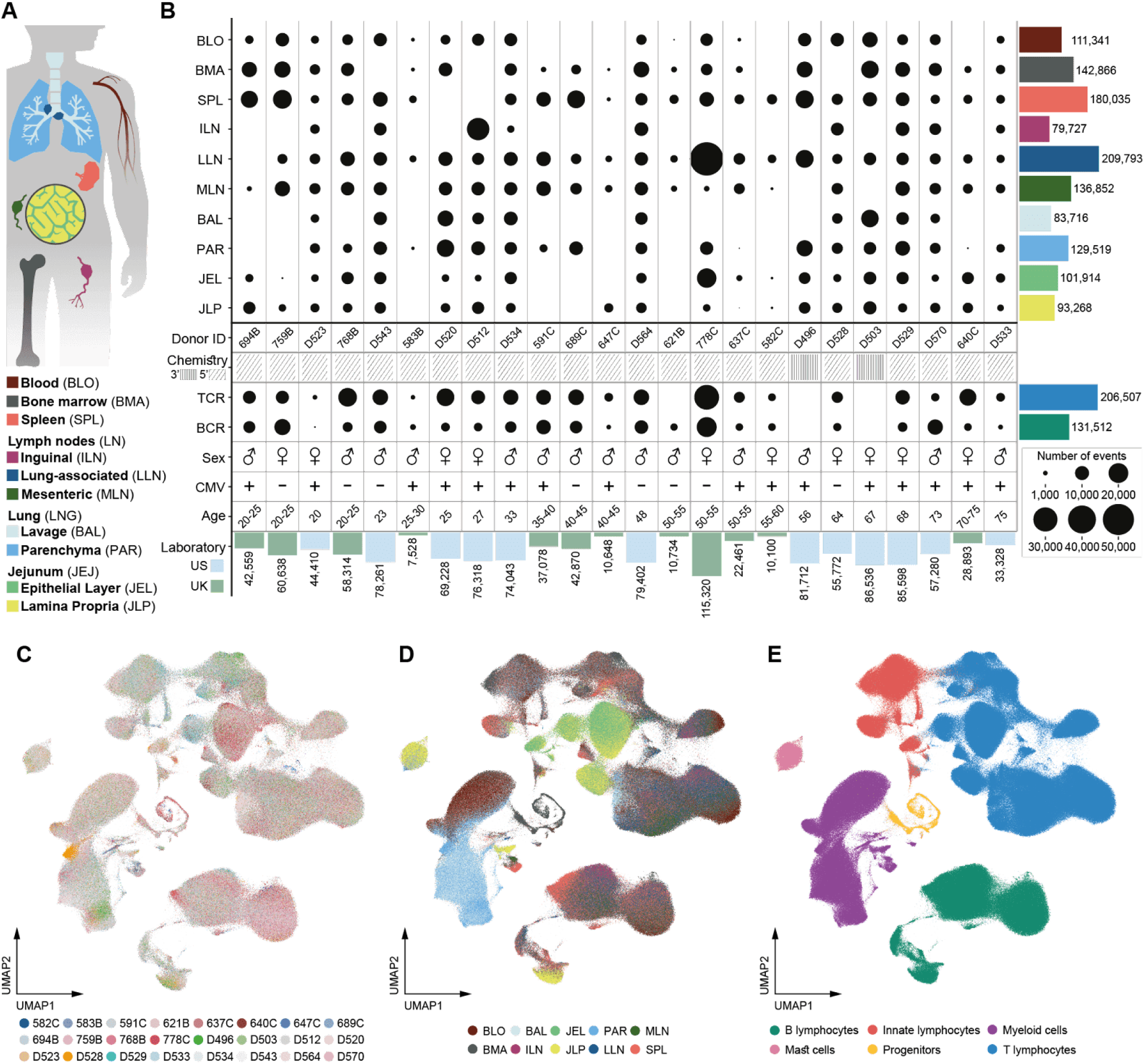
A multi-lineage human immune cell atlas across tissues and age. **(A)** Schematic of ten tissue sites used in analysis. **(B)** Dotplot of cells by donor and tissue site showing for each donor the sequencing chemistry, sex, CMV status, age, and whether the respective samples were obtained and processed in NY or UK. The scRNAseq data from 1,269,031 events are visualized as UMAP embeddings colored by donor. Bars in the bottom of the panels depict the overall number of cells profiled from each donor. Bars on the right of the panels depict the overall number of cells in each tissue across all donors. **(C)**, by tissue site **(D)**,or by major lineage as determined by the MMoCHi classifier **(E)**.

To integrate data from the ten key tissue sites and 24 donors we leveraged multi-resolution variational inference (MrVI). Designed particularly for cohort studies, MrVI provides two levels of integration: one that primarily reflects variation between cell states (allowing for unified annotation of cell states in all samples) and another accounting for differences between samples (allowing for comparative analysis) ^17^. The integrated dataset was visualized using uniform manifold approximant and projection (UMAP) ^18^, showing similar results across US and UK donors (**Fig. 1C, Extended data Fig. 1**). When visualized by tissue of origin, immune cells from blood, BMA, and LN integrated similarly, while cells from lungs, jejunum, and to a lesser extent, spleen localized by tissue (**Fig. 1D**). For immune cell subset annotation, we used the MMoCHi (MultiModal Classifier Hierarchy) algorithm ^19^ that leverages both gene and cell surface protein expression to hierarchically classify cells into predefined categories (**Supplementary Fig 1, Supplementary Tables 3, 4**). MMoCHi identified six major immune cell lineages: T lymphocytes, innate lymphocytes, B lymphocytes, myeloid cells, mast cells, and progenitors, each of which clustered distinctly and exhibited various levels of integration by tissue (**Fig. 1D,E**).

Using MMoCHi, we further defined thirteen T cell subsets, five NK/ILC subsets, six B cell subsets and seven myeloid subsets (**Fig. 2**). Manual curation was also performed for greater depth of annotation, resulting in 74 cell types and states (**Extended data Fig. 2, Supplementary Fig. 2**). The T cell compartment (610,429 cells) was classified into relatively rare γδ T cells, which develop early in ontogeny, and predominant αβ T cells consisting of CD4^+^ and CD8^+^ lineages subdivided into naïve, terminal effector (TEMRA), and memory – effector-memory (TEM), central memory (TCM), and tissue resident memory (TRM) – subsets, along with CD4^+^ regulatory T cells (Treg) defined based on surface markers and lineage defining genes ^16,20,21^ (**Fig. 2A, B**). Surface markers were used to enable the identification of T cell subsets that were not readily resolved by scRNA-seq alone^11,12^. For example, CD45RA surface expression was used to distinguish naive (CD45RA^+^) from TCM (CD45RA^-^), as each exhibit lymphoid homing profiles (CD62L, *CCR7*), and TEM (CD45RA^-^) from TEMRA (CD45RA^+^) cells, as both have effector-like profiles. In addition, TRM were distinguished from the TEM subset based on surface expression of core markers CD69, CD103, and CD49a ^16^ (**Supplementary Table 3**). TEM and TEMRA populations could be further subdivided into functional subsets based on expression of genes encoding cytolytic mediators (*GZMB*, *GZMK*) and the cytokine IL-17 (**Extended data Fig. 2** and **Supplementary Fig. 2**).

**Figure 2:**
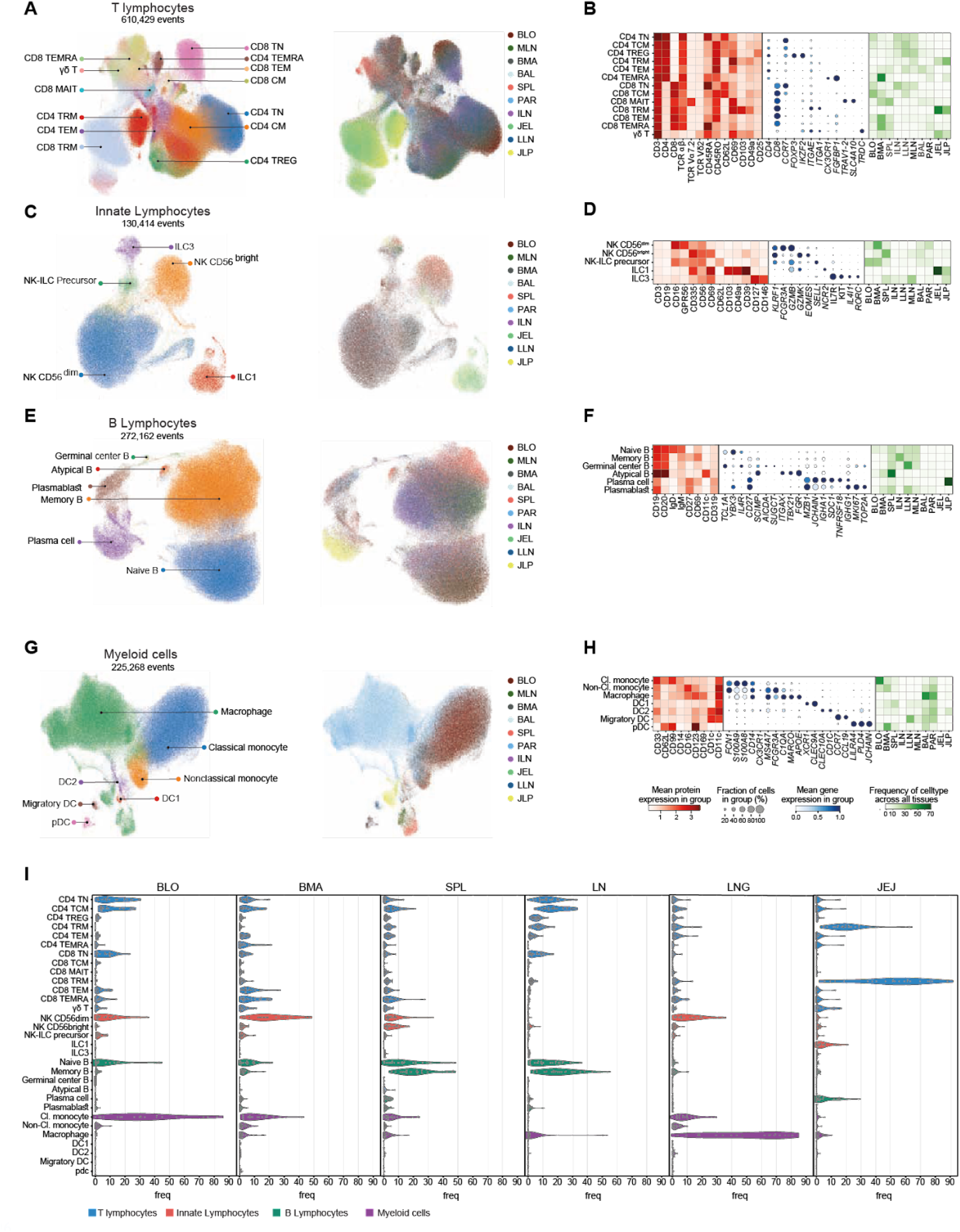
Heterogeneity and tissue distribution of immune cell subsets within the major lineages. Annotation of the T cell **(A,B**), NK\ILC **(C,D**), B cell (**E,F**), and myeloid cell (**G,H**) subsets was done using MMoCHi. (**A, C, E, G)** The annotated populations for each lineage are projected onto the UMAPs (left) and colored by tissue (right). (**B, D, G, H)** Next to each UMAP embedding is the protein (red matrix plot) and genes (blue dotplot) expression heatmap used to define each population and their relative distribution in each tissue (green heatmap; each row sums to 100%) calculated after downsampling to 79,666 cells (number of cells in the least abundant tissue - ILN) to adjust for the variability in cell number (see Abbreviations section for definitions of cell type labels and tissue labels). **(I)** Immune cell composition by tissue. The cell type composition is displayed as a violin plot where each dot is the frequency of a subset within each donor (frequencies sum to 100% for each donor). Abbreviations: BLD, blood; BMA, bone marrow; SPL, spleen; LN, lymph node: LNG, lung and bronchial alveolar lavage; JEJ, jejunum.

When visualized by tissue, naïve and TCM subsets were distributed in blood, BMA and lymphoid organs, TRM were predominant in the jejunum and found in lung and LN, and Treg were enriched in LN (**Fig. 2B**), consistent with previous flow cytometry analysis ^9,12,22^. Mucosal-associated invariant T (MAIT) cells distinguished by TCR expression, CD161 and other markers^23^ were enriched in spleen, BMA, and LNG (**Fig. 2A,B**). T cell receptor (TCR) clonal analysis provided additional correlative evidence for T cell subset delineation and tissue distribution (**Supplementary Fig. 3**). In particular, CD8^+^ T cells exhibited greater clonal expansion compared to CD4^+^ T cells, the TEMRA subset had the highest clonality across sites followed by TEM cells, and tissues with predominant memory T cells had the highest clonality compared to LN which contain naive populations with high diversity (**Supplementary Fig. 3**). These findings were consistent with previous TCR analysis of tissue subsets ^12,24^.

Innate lymphocytes (130,414 cells) across tissues consisted mainly of the CD56^dim^ mature NK subset, characterized by surface expression of CD16 and genes associated with cytolytic function (*KLRF1*, *GZMB*), along with the less abundant CD56^bright^ (immature) NK cells with lower CD16 expression (**Fig. 2C,D**). Innate lymphoid cells (ILC) including ILC1 (CD16^-^*NCR2*^+^*IL7R*^-^) and ILC3 (*IL7R*^+^*KIT*^+^*RORC*^+^)^25^ subsets were also detected, but at lower frequencies (**Fig. 2C,D**). Across tissues, CD56^dim^ cells were present mostly in blood, BMA, spleen, and lungs, while CD56^bright^ cells were found in most tissues and enriched in the spleen (**Fig. 2C,D)**. ILC1, which expressed high levels of the tissue residency markers CD69, CD49a and CD103, were found predominantly in the jejunum, while ILC3s were mainly in LN, spleen, and mucosal sites (**Fig. 2C,D**). Putative NK-ILC precursors, resembling both CD56^bright^ NK and ILC3s (CD127^+^CD44^+^*GPR183*^+^) were found mainly in blood, BMA, lung and spleen (**Fig. 2C,D** and **Supplementary Fig. 2**), while a population with characteristics of ILC1 and ILC3 (ILC1-3 “Transitional”, *NCR2*^+^CD127^+^) was exclusive to jejunum (**Extended data Fig. 2** and **Supplementary Fig. 2**)

The B cell compartment (272,162 cells) (**Fig. 2E,F**) was classified into six major subsets including: naive B cells (IgD^+^), memory B cells (CD27^+^), germinal center (GC) B cells that express *AICDA* (activation-induced adenosine deaminase (AID) for immunoglobulin class switch recombination ^26^), plasma cells that express *SDC1* (CD138) and highly express immunoglobulin genes, and plasmablasts that express proliferation-associated genes (*MKI67*, *TOP2A*). A subset of “Atypical B cells” (CD27^-^IgD^-^) which express *TBX21* encoding the transcription factor T-BET ^27,28^ was identified at low frequencies across multiple sites (**Fig. 2E,F, Extended data Fig. 2**). Manual annotation revealed an additional low frequency population of pro-B cells expressing *MME*, *SOX4* and CD38 restricted to the bone marrow (**Extended data Fig. 2** and **Supplementary Fig. 2**). Overall, B cells were found mostly in lymphoid organs and blood; naïve B cells were in blood, spleen, and LN, and memory B cells were predominantly distributed in spleen and LN. A proportion of these lymphoid memory B cells expressed CD69, denoting tissue resident B cells ^29^. Plasma cells were enriched in the jejunum lamina propria while plasmablasts were enriched in lymphoid organs (**Fig. 2E,F**). B cell receptor (BCR) analysis provided additional confirmation and insights into subsets and tissue distribution; plasmablasts exhibited the highest clonal expansion across lymphoid sites, while memory B and plasma (but not naive) cells expressed mutated BCR (**Supplementary Fig. 4A-G**). Across sites, jejunum was enriched with plasma cells expressing IgA, while blood contained fewer memory B cells and lower mutation frequencies compared to all tissues examined (**Supplementary Fig. 4H-J**)

Myeloid-lineage cells (225,268 cells) comprised of macrophages, classical and non-classical monocytes, and dendritic cell (DC) subsets, including DC1 (*CLEC9A*^+^), DC2 (*CLEC10A*^+^), migratory DC (*CCR7*^+^), and Plasmacytoid DC (pDC; CD123^+^*LILRA4*^+^) (**Fig. 2G,H**). Classical monocytes were enriched mostly in blood while non-classical monocytes were also found in BMA and lung. pDCs were enriched in the bone marrow, while other DC subsets were found in the lung, jejunum, and LN (**Fig. 2G,H**). Macrophages were found predominantly in the lungs, but were also present at low frequencies in the jejunum, spleen, and LN (**Fig. 2H, Extended data Fig. 2**). By manual annotation, macrophages are further subdivided, into *MARCO*^+^ macrophages enriched in the lung, *SPIC*^+^ macrophages enriched in the spleen and *SDS*^+^ macrophages that are found predominantly in the jejunum (**Extended data Fig. 2** and **Supplementary Fig. 2**). Collectively, this comprehensive annotated map of immune cells across major lineages and tissues using multimodal profiling can serve as a reference for annotations, embeddings, and other types analyzes to enable future studies; e.g., using pre-trained models in the popV framework ^30^ (**Supplementary Fig. 5**).

Collating all of the lineage and subset annotations within and across tissues reveals tissue-specific immune cell compositions with some lineages showing varying proportions between donors (**Fig. 2I, Extended data Fig. 2, Supplementary Fig. 6**). Blood, BMA and spleen contain a similar complement of immune cell subsets with some site specific differences; BMA contained fewer naïve T cells, additional memory T cell subsets, more NK cells and fewer monocytes relative to blood, while spleen contained more memory B cells, NK cell subsets, and lower monocyte proportions compared to blood. Moreover, LN contained increased naïve, TCM, TRM, memory B cells, and macrophages and fewer monocytes compared to spleen and BMA. Lungs and jejunum were most distinct from lymphoid organs and each other; lung contained the greatest proportion of macrophages, low frequencies of B cells and multiple memory T cell subsets, while intestines contained largely TRM populations, ILC1, macrophages and low frequencies of B cells. These analyses reveal a tissue intrinsic nature of immune cell composition for all immune cell lineages.

### Tissue is a major determinant of immune cell identity and function

To understand how tissue localization influences the transcriptional state of immune cells, we used MrVI to calculate similarity between the ten tissue sites for each of the major immune cell lineages. Hierarchical clustering of the average MrVI distances across lineages revealed high levels of transcriptional similarity within three tissue groups: jejunum (JEL, JLP), lymph nodes (MLN, ILN, LLN) and lung (BAL, PAR), while the remaining sites (blood, BMA, SPL) clustered separately (**Extended data Fig. 3A**). We then performed differential expression (DE) analysis, separately for every major lineage (CD8^+^ T, CD4^+^ T, Myeloid, NK/ILC and B cells), comparing each tissue group against the remaining five tissue groups using a pseudo-bulk linear mixed model approach that accounts for the major nuisance covariates in our data (e.g., donor ID, sex, donation site, CMV status)(**Supplementary Table 5**) ^31^. The sets of genes from each tissue group that were significantly DE in three or more of the major immune lineages were then merged and hierarchically clustered, resulting in ten gene expression groups and five lineage-tissue groups (**Fig. 3A**). This approach revealed how functional gene profiles are distributed across lineages and tissues.

**Figure 3:**
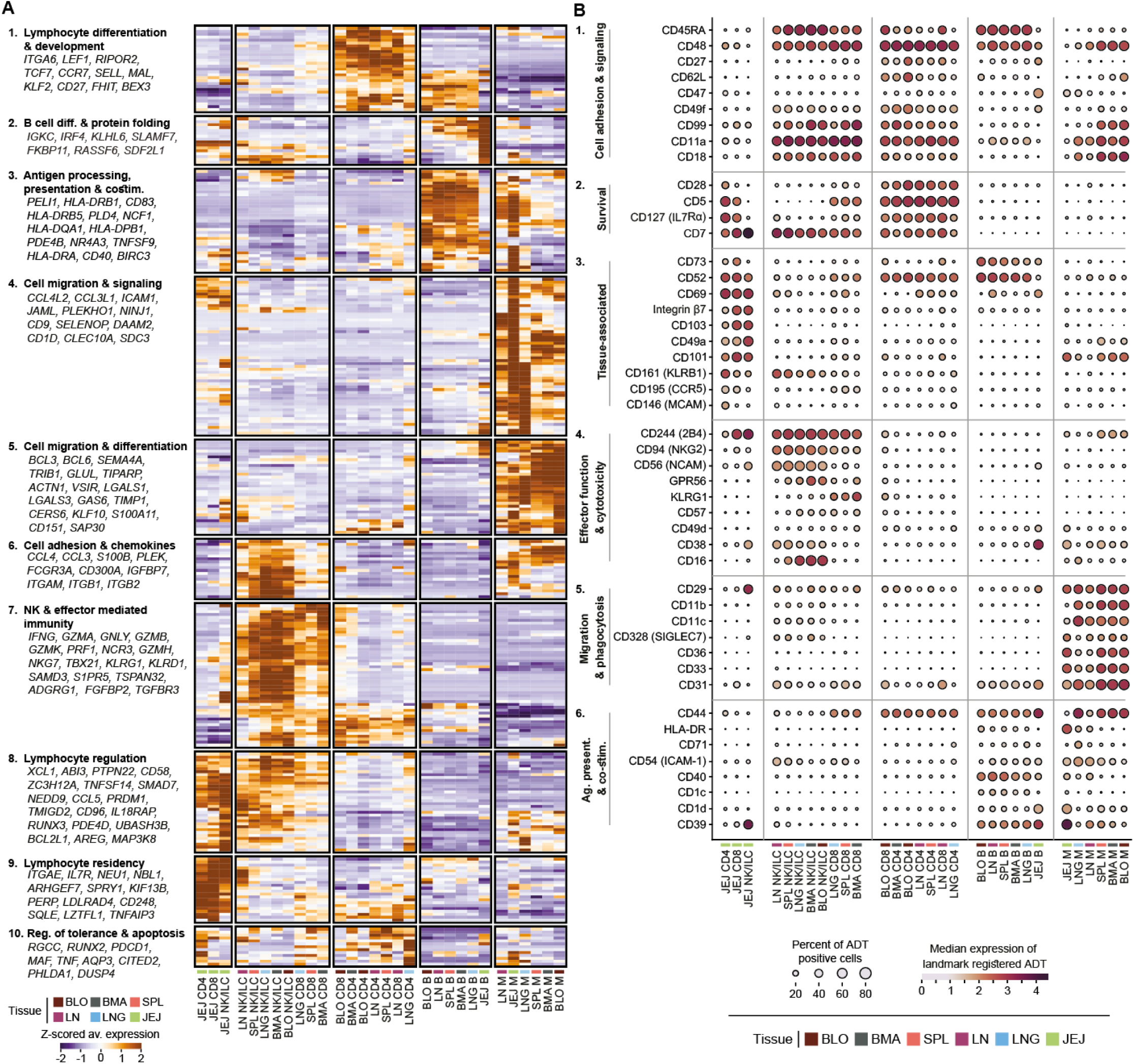
Tissue is a major determinant for immune cell composition and identity. **(A)** Heatmap depicting the expression (Z-score of average expression across cells) of tissue-associated genes. Included are genes that were detected (adj. p-value<0.05, LFC>1) in at least three lineages in at least one of the tissue groups (compared to all other tissue groups). **(B)** Dot plot of significant DE proteins (adj. p-value<0.05, LFC>0.585). Included are proteins that were detected as DE in at least two lineages in at least one of the tissue groups. As the protein DE analysis is conducted separately for each donor, we also require the displayed DE results to hold in at least 65% of donors.

Certain functional profiles were enriched in specific cell lineages and found across multiple tissue sites (**Fig. 3A**); for example, myeloid cells exhibited specific cell migration and differentiation profiles across all tissues (cluster #4, 5). Other clusters were shared across distinct lineages such as the antigen presentation and costimulatory programs (cluster #3) shared between B and myeloid cells in blood, lymphoid organs, and lung, but lowly expressed the B cells of the jejunum. Another example is the chemokine and cell adhesion cluster (cluster #6) shared between innate lymphocytes, myeloid cells and to a lesser extent CD8^+^ T cells (**Fig. 3A**). Furthermore, a cluster of genes associated with stem cell-like features (*ITGA6, LEF1, TCF7*) and lymphoid trafficking (*SELL, CCR7*) was enriched within T cells in the blood and LN, and a subset of the genes defining this signature was enriched in B cells in blood, lymphoid organs, and lungs(Cluster #1). Conversely, genes associated with effector function (*IFNG, TBX21, FCGR3A*), and cytotoxicity (*GZMB, GZMK, PRF1, GNLY*) (Cluster #7) were shared between CD8^+^ T cells, NK/ILC across all tissues except the jejunum, and a subset of genes within this cluster were present in CD4^+^ T cells in the BMA, spleen, and lungs. Finally, there were several gene expression clusters markedly enriched in NK/ILC and T cells in the jejunum, including genes associated with tissue residence (*ITGAE*, *SPRY1*, *CD248*) (Cluster #9), TRM differentiation (*PRDM1*, *RUNX3*)^32,33^, tissue repair (*AREG*)^34^, and control of apoptosis and signaling (*BCL2L1*, *PTPN22*) (Cluster #8, 10) (**Fig. 3A)**.

To further investigate these functional signatures across lineages and tissues, we performed differential expression on the surface protein data between tissue sites for each individual donor (**Supplementary Table 6**). Protein abundances were then hierarchically clustered into six major groups, revealing functional signatures similar to the gene expression analysis. Proteins associated with migration and phagocytosis (CD11b, CD11c, CD36; cluster #5) were expressed by myeloid cells in all tissues. Furthermore, proteins associated with antigen presentation (e.g. HLA-DR, CD40, CD1d; cluster #6) were shared between myeloid and B cells in lymphoid sites and the lungs, but less so in the gut B cells (**Fig. 3B**, Clusters #5, 6). We also found a stem-like, lymphoid homing cluster which included expression of CD62L, along with integrins CD49f, CD18, and CD11a by T and NK cells in blood, lungs, and lymphoid organs (**Fig. 3B**, Cluster #1). The effector and cytotoxic cluster (#4) expressing CD16, KLRG1, CD244, GPR56, and CD94 was identified in CD8^+^ T cells and NK/ILC from multiple sites. Finally, molecules associated with tissue residence including CD103, CD49a, CD101, integrin β7, and CD69 were expressed by T and NK/ILCs, and were highly enriched in the jejunum (**Fig. 3B**, Cluster #3). Together our differential expression analysis validated and extended key signature for immune cell functional states across lineages and tissues.

To specifically interrogate the overall role of tissue in immune cell profiles compared to the more readily profiled and accessible blood, we performed compositional and differential expression analysis of each major lineage in blood versus each tissue. We identified compositional differences between the blood and each tissue site as well as differentially expressed genes across multiple sites for each lineage (**Extended data Fig. 3B-C, Supplementary Fig. 6H**). We found that localization in tissues is accompanied by broad changes in expression of genes associated with metabolism, DNA repair, antigen presentation, cell migration, and post-translational protein modifications. These results suggest that tissue localization may have generalized effects on immune cell functions which are further modified and honed for residence in specific sites, and thus obscured in studies of PBMCs.

### Tissue-specific transcriptional programs across immune lineages

In order to assess whether the tissue imparted distinct gene signatures across lineages, we implemented consensus single cell hierarchical Poisson Factorization (scHPF)^11,35^, a Bayesian factorization algorithm that identifies the major co-expression patterns (“factors”) in the data, rather than considering each gene in isolation (as we have done for DE). We applied scHPF to each major immune cell lineage in separate models with balanced cell input across tissues (**Supplementary Table 7**), which allowed us to identify a variety of transcriptional programs. We next identified which of the factors were enriched in individual tissues, and hierarchically clustered these factors to reveal distinct groups of gene expression programs or “modules” that were strongly associated with one or more tissues, many of which spanned multiple cell lineages (**Fig. 4A).**

**Figure 4:**
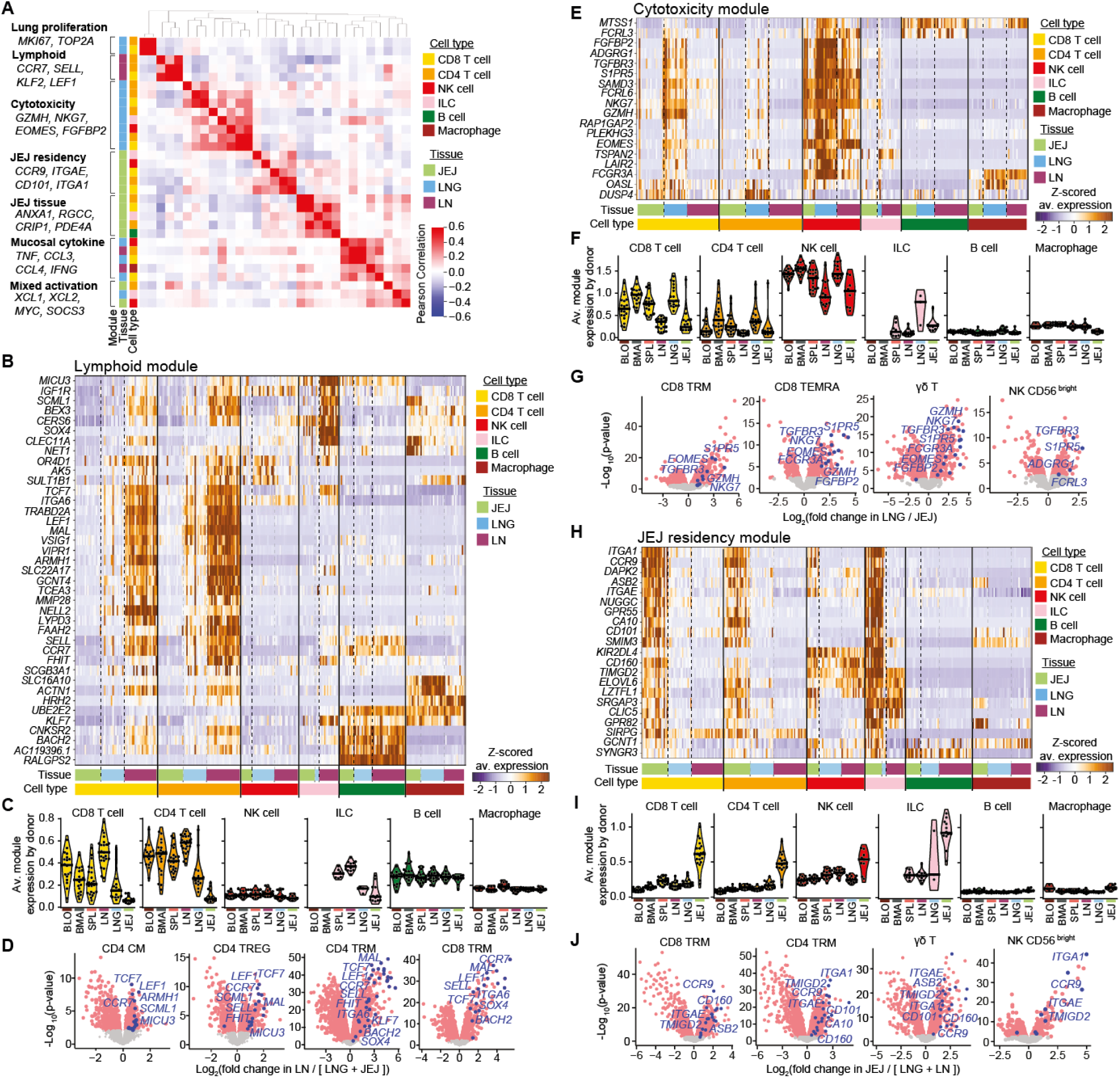
Co-expression programs across immune subsets in tissues. Consensus scHPF was used to factorize gene expression profiles of six immune lineages across three tissues to generate a list of tissue-specific factors. **(A)** Correlogram showing hierarchically-clustered Pearson correlation values between factor gene scores. Seven gene expression modules containing multiple factors with similar transcriptional signatures were identified. The cell type model and tissue enrichment for each of the factors within modules are indicated, with selected top genes within each module listed on the right. **(B)** Heatmap showing z-scored average expression of genes within the Lymphoid module across immune cell subsets and tissues used by scHPF. **(C)** Violin plots showing averaged expression of Lymphoid module genes within a donor across immune cell subset and expanded tissue groupings. **(D)** Volcano plots showing differential expressed transcripts (pink) within selected immune cell subsets across indicated tissues. Genes from the Lymphoid module are indicated in blue. **(E), (F), (G)** Same as **(B), (C)** and **(D)**, but for genes in the Cytotoxicity module. **(H),(I), (J)**, Same as **(B), (C)** and **(D)**, but for genes in the Residency module.

This factorization approach identified three tissue-specific gene expression modules that were manifested across lineages and subsets (**Figs. 4B-J**). Furthermore, as we show by examining module expression within finer-grained immune cell subsets between tissue types by differential expression (**Supplementary Table 8**), these tissue-specific modules are independent of subset composition. The lymphoid module included genes associated with quiescence (*KLF2*, *LEF1, KLF7*) and lymphoid homing (*CCR7*, *SELL, ITGA6*) ^36,37^, which were enriched in T, B and NK/ILC in LNs and not present in mucosal sites (**Fig. 4B, C**). Importantly, multiple subsets of T cells, including CD4^+^TCM, CD4^+^Treg, CD4^+^TRM, and CD8^+^TRM, all showed significantly increased expression of lymphoid module genes in the LN compared to mucosal sites (**Fig. 4D**). Second, a “cytotoxicity module” which consisted of multiple effector- and cytotoxicity-associated genes (*GZMH, EOMES, NKG7, FGFBP2, ADGRG1*(GPR56), *FCGR3A*), was highly expressed by T cells, NK cells and ILCs in the lung compared to LN and gut, and also enriched in blood, bone marrow and spleen (**Fig. 4A,E, F**). Moreover, CD8^+^TRM, CD8^+^TEMRA, γδ T cells and CD56^bright^ NK cells all showed significant enrichment of the cytotoxicity module genes in the lungs when compared to their counterparts in the gut (**Fig. 4G**), further validating the identification of a tissue-driven gene expression program. Lastly, a gut residency module contained genes specific to the jejunum and associated with tissue residency (*CCR9*, *ITGAE*, *CD101*, *CD160*)^12,38^ for CD4^+^ and CD8^+^ T cells, NK cells and ILCs, while absent in other sites (**Fig. 4H,I**). This module also showed increased expression in CD4^+^TRM, CD8^+^TRM, γδ T cells and CD56^bright^ NK cells in the jejunum compared to the lung and LN (**Fig. 4J**). Taken together, these analyses establish that tissue has major effects not only on immune cell composition, but also immune cell transcriptional programs.

### Immune cell aging signatures manifested by specific lineages, subsets, and sites

The age range of the donors spanning six decades of adult life (20–75 years) enabled investigation of age-associated effects across immune lineages and anatomic sites. A global (pseudo-bulk) analysis of variation reveals that while the tissue environment is the dominant factor in the molecular profiles across lineages, aging still explains a fraction of the variance, with some of the most affected genes representing key pathways of immune activation and migration (**Extended data Fig. 5A**). Overall, immune cell composition within each site was maintained over adult age; however, there were decreased frequencies of naïve CD8^+^ T cells and a decline in subpopulations within naïve B cells, and plasmablasts with age, mostly manifested in blood and LNs (**Extended data Fig. 5B, Supplementary Fig. 7**).

**Figure 5:**
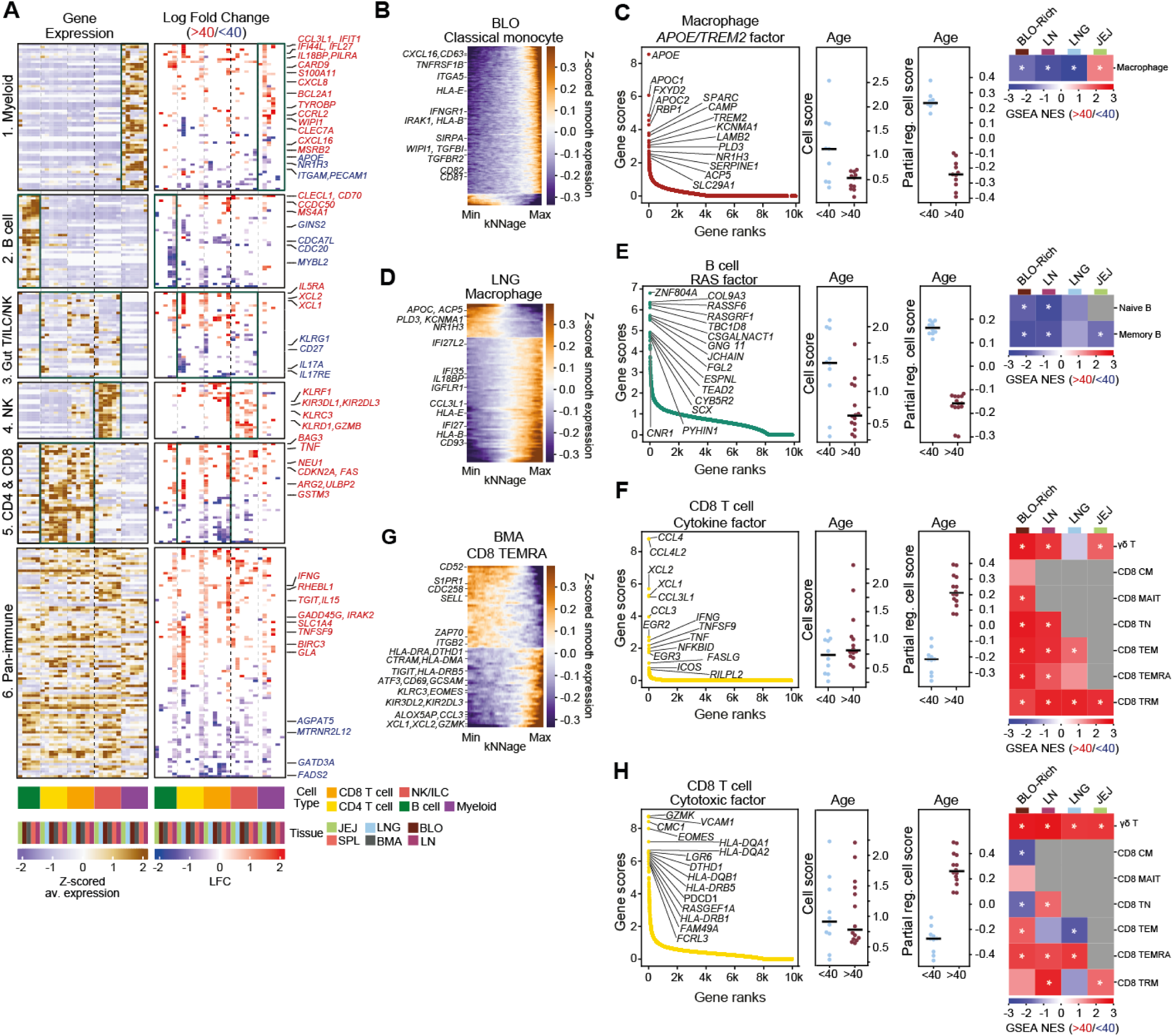
Aging signatures across immune subsets and tissue locations. **(A)** Heatmap depicting age-dependent genes across at least two lineages or two tissues. Clustering by mean expression levels in each combination of tissue-cell lineage (left panel) revealed six groups, highlighting up-regulated (blue) or downregulated (red) genes associated with age (right panel). Fold changes are only displayed for tissue-cell lineage combinations in which the respective gene is expressed at sufficient levels (see Methods) and significantly changes with age (adj p<0.05, |LFC|>0.1). Black lines indicate the most relevant (highest expression) combination of tissue and cell lineage in each gene group. **(B)** kNNage analysis toidentify age-dependent genes in blood monocytes. For this analysis we first assigned each cell with a smooth age coefficient by averaging the donor ages of its 50 nearest neighbors in mrVI latent space. Expression of genes that significantly correlated with these age coefficients (empirical p<0.1, |R|>0.1) is displayed with cells ordered by their age coefficients. Displayed values are Z-scored and smoothed with a sliding window of 5% of cells. **(C)** Dot plot indicating rank and gene scores for the top genes in the *APOE/TREM2* factor derived from the consensus scHPF macrophage model is shown on the left. Factor cell scores and partially regressed cell scores by donor age group are shown in the middle. Heatmap of normalized enrichment scores (NES) by pre-ranked GSEA in the discrete differential expression analysis by cell subset using top 100 ranked genes in the factor as the gene set. **(D)** Same as **(B)** but for macrophages from the lung. **(E)** Same as **(C)** but for the B cell *RAS* Factor identified by consensus scHPF. **(F)** Same as**(C)** but for the CD8^+^ T cell Cytokine Factor identified by consensus scHPF. **(G)** Same as **(B)** but for CD8^+^ TEMRA from bone marrow tissues. **(H)** Same as **(C)** but for the CD8^+^ T cell Cytotoxic Factor identified by consensus scHPF.

To identify specific gene expression programs that change with age within and across tissues, we applied two complementary approaches. First, we performed a covariate-corrected DE analysis separately for each major immune lineage across tissue groupings and relied on both discrete – comparing pseudo-bulk profiles in younger (< 40 years) versus older (> 40 years) individuals – and continuous tests – single-cell level, considering all ages (**Supplementary Tables 9, 10, and Supplementary Fig. 8**). Second, we generated scHPF models separately for each major lineage with balanced representation for each donor (**Supplementary Table 11**), regressing out covariates to identify expression programs that change with age – comparing normalized scHPF factor scores in younger versus older individuals. Our analysis revealed different classes of genes that are associated with age, including those associated with specific lineages, those associated with specific sites, and a “global aging signature” found across many lineages and sites (**Fig. 5**).

Among myeloid cells, genes encoding pro-inflammatory cytokines, chemokines, and interferon-regulated genes were increased with age in the lung (e.g., *IFIT1/27/44L*, *IL18BP*, *CCRL2*, and *CCL3L1*) and blood (*CXCL16*, *S100A11*, *SLC11A1*) (**Fig. 5A**, cluster #1). This age-associated increase in pro-inflammatory genes was also observed in classical monocytes from the blood (*CXCL16*, *IFNGR1*, *IRAK1*, *TNFRSF1B*) (**Fig. 5B)**, and in monocytes and macrophages from the lung (*IFIT3/44L*, *IL18BP, MARCO, IL1RN,* **Supplementary Fig. 9A-B**). Conversely, lung macrophages exhibited an age-related decrease in expression of genes encoding apolipoproteins (*APOC1*, *APOC2* and *APOE*), and *TREM2* – a triggering receptor expressed on myeloid cells ^39^ that binds the lipid binding protein, ApoE, and facilitates macrophage functions, such as phagocytosis and chemotaxis, and induces metabolic changes ^40^ (**Fig. 5A,C,D**). This *APOE-TREM2* containing module was also identified by scHPF analysis as being decreased in older compared to younger individuals (using the same age cutoff) within myeloid cells in blood-rich tissues (BMA, spleen, blood), LNs, and lungs, but not in the gut (**Fig. 5C**). We validated the age-associated decrease in expression of the *APOE-TREM2* gene module in an independent dataset from human lungs ^41^ **(Extended data Fig 5C**). While extensive studies have revealed a significant association of *APOE-TREM2* expression in microglia with neurodegeneration in Alzheimer’s disease and with macrophages in cardiometabolic disorders in mice and humans ^40,42,43^, the decreased expression with age in mucosal and lymphoid sites identified here, appears to to be a normal feature of aging.

For B cells, there were age-associated increases in expression of genes associated with T-B cell interactions, especially in the spleen and bone marrow, including *CD70* (encoding the ligand for CD27 expressed by memory B cells), *MS4A1* (CD20) and *CLECL1* ^44^, along with decreased expression of genes involved in cell cycle control (*CDC20, MYBL2*) (**Fig. 5A**, cluster #2). We further identified by scHPF an age-associated decrease in the expression of a gene module associated with RAS signaling (*RASA4B*, *RASGRF1*, *PLAAT3*, *GAB2*) across multiple sites and subsets (**Fig. 5E**). To test the reproducibility of this result, we confirmed that the expression of RAS genes is enriched in bone marrow-derived B cells from younger compared to older donors in an independent dataset^45^ (**Extended Data Fig. 5C**). The RAS pathway is coupled to B cell receptor-mediated signaling ^46^ and has specific roles in memory B cell activation, differentiation and class switching ^47^. We also found an age-associated increase in *CD99* within lymph node memory B cells (**Supplementary Table 10**), a surface molecule which signals through the RAS pathway and regulates cell adhesion, survival, and immune cell interactions ^48^. Our results therefore suggest alterations in memory B cell signaling and differentiation with age.

Among T and NK cells, our analysis identified several aging programs, using DE and scHPF, which are primarily associated with pro-inflammatory and cytotoxic function. One gene module we identified, comprised of multiple pro-inflammatory cytokines and chemokines (e.g., *CCL3*, *CCL4*, *XCL1*, *CXCL2*, *IFNG*, *TNF*) and was increased with age in multiple CD8**^+^**subsets, most prominently TRM, across all tissues (**Fig. 5A, F**). A second gene module consisting of cytotoxicity-associated genes (*KLRC3*, *KLRD1*, *KLRF1*, and *GZMB*) increased with age, predominantly in NK at lung and blood-rich sites (**Fig. 5A**, cluster #4, **Supplementary Fig. 9A**). Finally, scHPF analysis identified an age-associated increase in a gene expression program containing cytotoxicity and activation-associated genes (*GZMK*, *EOMES*, *PDCD1*, *HLA-DR*) manifested by γδ T cells in all sites and CD8^+^TEMRA in all sites except the gut (**Fig 5H**). Compositional analysis confirms that there is an increased frequency of *GZMK*-expressing CD8^+^TEMRA cells in older donors (**Extended Data Fig 5D**), consistent with results in aged mice ^49^. By continuous analysis specific to CD8^+^TEMRA in BMA we further observe increased expression of *GZMK* along with multiple pro-inflammatory chemokines, additional cytotoxicity genes (*KIR3DL2*, *KIR2DL3*) and activation markers (*CD69*, *HLA-DR*, *TIGIT*) (**Fig. 5G**). Finally, TCR analysis shows that these *GZMK*^+^CD8^+^TEMRA exhibited increased clonal expansion in older donors (**Extended data Fig. 5E**). Taken together, these results suggest gradual alteration in the activation and effector function of the TEMRA population with age.

The differential expression analysis identified a cluster of genes that exhibited age-related changes across most lineages (**Fig. 5A**, cluster #6). This putative global immune aging signature included increased expression of genes associated with IFN signaling (*IFITM1*, *IFIT2*), mTOR pathways, (*GLA*, *FADS2*, *SLC1A4*, and *RHEBL1* ^50^), and pro-inflammatory responses (*PTGER2*, *IRAK2*, *RTP4*, and *TNFSF9*), as well as a marked decrease in expression of multiple genes related to mitochondrial metabolism (*GATD3A*, *AGPAT5*, and *MTRNR2L12)* ^51^. These global changes across lineages and tissues are consistent with increased inflammation or “inflammaging” being a major part of the aging process resulting in altered metabolism, increased disease susceptibility, and organ damage ^52^.

### Tissue-specific influence of age on the continuum of T helper cell states

While the above analysis examined the effects of age on annotated cell subsets, it is conceivable that some aging effects may be restricted to parts of an annotated subset and/or span several subsets. This is especially the case for T helper (CD4^+^) cells, which are highly abundant in our data, shared across all tissue sites, and often display a phenotypic continuum rather than separation into well-distinguished subsets ^53,54^. For an annotation-independent analysis of aging in CD4^+^ cells, we leveraged a per-cell estimation of the pertaining effects using counterfactual analysis with MrVI, which considers each cell separately and controls for unwanted covariates (see Methods). This analysis revealed age-related changes for CD4^+^ T cells in the lung, gut, and lymph nodes that were specific to these sites or manifested differently in other sites (**Fig. 6**) (see complete set of results, for every tissue group and subset in **Extended Report 1**).

**Figure 6.**
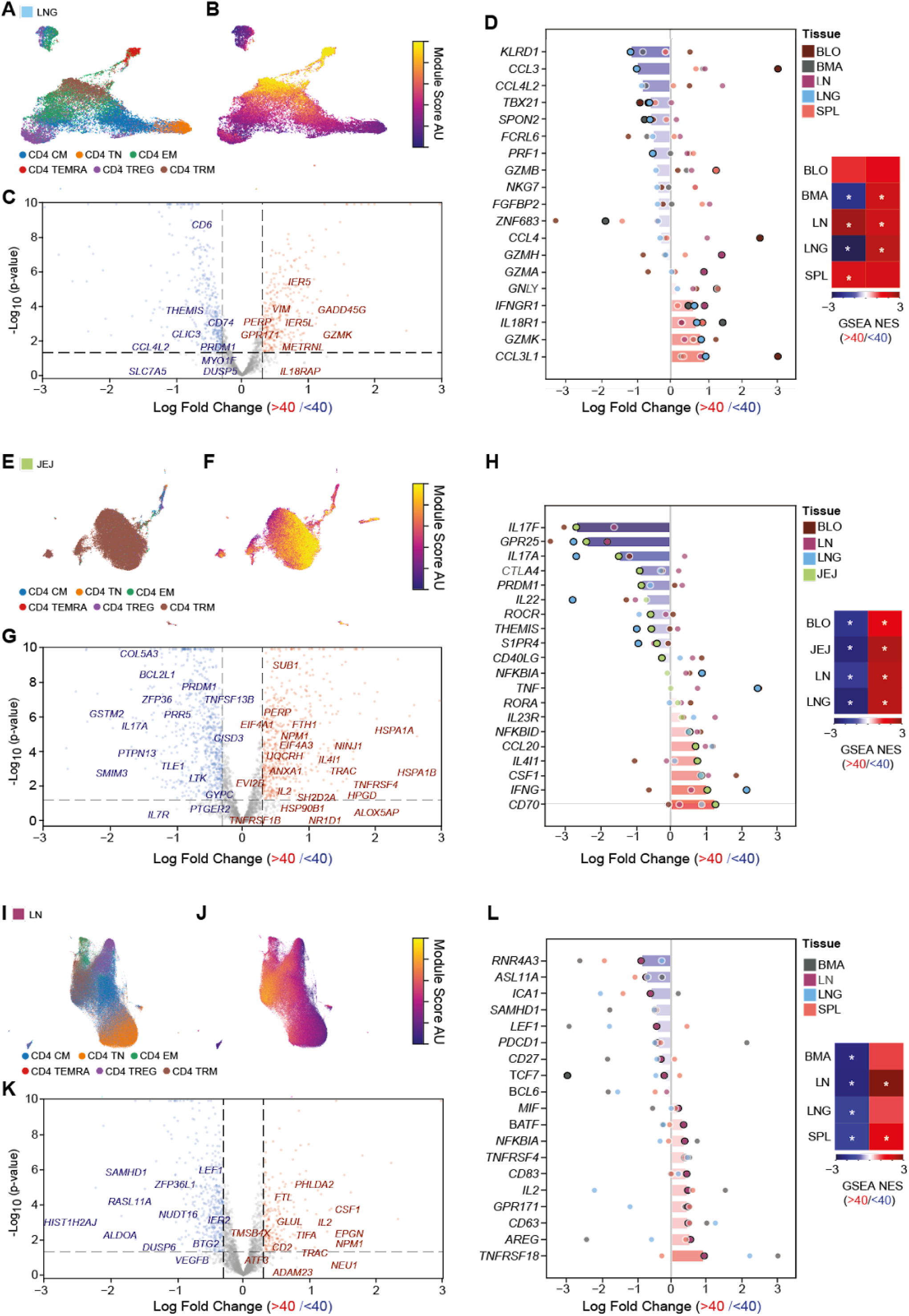
Tissue-dependent signature of aging in CD4^±^ T cells. MrVI analysis was performed to evaluate age-related effects (LFC) for each gene and in every cell (comparing donors over 40 to under 40 years old). Spectral biclustering of the LFC cell x gene matrix was used to identify gene signatures (i.e., sets of genes with similar aging effects) and the subpopulations of cells in which these effects are manifested (signature-positive cells)**.(A)** UMAP embedding of CD4^+^ T cell-types in the lung. **(B)** UMAP with cells colored by the lung cytotoxicity signature score. Scores are computed by summing the inferred LFCs of signature genes that are upregulated in old individuals minus the sum of LFCs of genes downregulated in old individuals. The respective signature-positive cells are highlighted by high signature scores and overlap with TRM and TEMRA. **(C)** The signature-positive cells were used for pseudo-bulk DE analysis (comparing donors over 40 to under 40 years old) and the top 50 genes from the gene signature (by estimated LFC) are displayed if their unadjusted P-value is below 0.1. **(D)** A classifier was trained to identify signature-positive cells in other tissues. Log-fold changes and p-values were computed using pseudo-bulk DE on these cells (as above). The bars denote the pseudo-bulk LFC in the lung. All organs are displayed by overlaying circles. Opaque and black circle signifies an unadjusted p-value below 0.05, while transparent and white circle means a p-value above 0.05. We present manually selected genes with known effector or activation functions, which either come from the signature or show similar trend by pseudo-bulk DE in the lung (panel C). GSEA was computed for manually-selected genes to explore their age dependency in other tissues. Separate analysis was conducted to genes that are up or down-regulated in the lung. Gene rankings for GSEA are based on pseudo-bulk DE in the respective tissue. NES scores are displayed and a star means significant with a p-value below 0.1. **(E, F, G,H)** and **(I, J, K, l)** similar to **(A, B,C, D)**. Tissues in **(D, H, I)** are displayed only if a sufficient number of signature-positive cells was detected. Note that differing amounts of signature-positive cells underlies missing significance of genes with high log-fold changes.

In the lung, we detected an age-related gene expression signature in CD4^+^ TEMRA cells marked by increased expression of certain cytolytic/effector-associated genes (*GZMK, IFNGR1, IL-18R*) concomitant with decreased expression of other genes associated with cytolytic effector function (*PRF1*, *TBX21*) and inflammatory chemokines (*CCL3*, *CCL4L2*)^55,56^ **(Fig. 6A-D)**. This pattern of age-related gene expression changes was distinct from CD8^+^TEMRA, in which age-associated increases in all of these effector-related genes was observed (**Fig. 5G-H**). We trained a classifier to identify similar sets of module-positive cells in each of the other tissues and compared the aging effects on these matched sets to those observed in the lung (Methods). We find that the upregulated effector genes in lung CD4^+^TEMRA were also upregulated with age in the corresponding populations in all other sites (e.g., blood, BMA, spleen, LN), while the downregulated genes were similarly observed in BMA but exhibited increased expression with age in the blood, spleen and LN **(Fig. 6D)**. Notably, these discordant genes are enriched in cytotoxic effector molecules, such as *PRF1*, *FGFBP2*, and *GZMH*. These results therefore point to age-associated changes in the regulation of cytotoxicity in a major subset of the CD4^+^ memory pool that is discordant between different tissue sites.

In the jejunum, where the majority of CD4^+^ T cells are TRM, this analysis highlights a subpopulation of cells with an age-related decline in genes involved in the Th17 pathway including *IL17A*, *IL17F,* and *IL22* and the associated transcription factor *RORC.* These changes are concomitant with increased expression of genes encoding proinflammatory cytokines (*IFNG*, *CSF1*) and activation of the NF-κβ pathway (*NFKBID*), which promotes inflammatory processes (**Fig. 6E-H**). The reduction in IL-17-associated genes coupled with increased *IFNG* has been associated with barrier dysfunction in the gut ^55^, suggesting a similar effect with aging. Because this CD4^+^TRM population is specific to the jejunum in our dataset, we investigated potential age-associated changes in CD4^+^*IL17A*^+^ cells across all sites where they are found at sufficient numbers in our datasets (i.e., blood, lung, and LN). This analysis revealed a similar age-associated decline in IL-17-associated genes coincident with an increase in proinflammatory cytokines (e.g., *CSF1*, *CCL20*) in all sites (**Fig. 6H**), suggesting a systemic loss of the Th17 signature with age.

A third major age-associated change for CD4^+^ T cells was found in LN, manifested by the loss of genes which define the T-follicular helper (Tfh) subset^57^ including *PDCD1* (encoding PD-1) and *ICA1* along with reduced expression of the stem-like factors *LEF1* and *TCF7*, and an increase in genes associated with T cell activation (*IL2*, *TNFRSF4*, *CD83*) and tissue repair (*AREG*, encoding amphiregulin)^34^ (**Fig. 6I-L**). Genes that we found downregulated in the LN show similar trends in the corresponding subpopulations in the lung, BMA, and spleen, while upregulated genes show a similar pattern mostly in the spleen while *AREG* upregulation is specific to LN (**Fig. 6L**). This age-related reduction in Tfh-associated genes across lymphoid sites indicates reduced help for B cell differentiation at the inductive sites for adaptive immunity, potentially accounting for the diminished humoral responses in older age ^58,59^.

## DISCUSSION

We present here a comprehensive, high dimensional, global analysis of the human immune system across tissues and ages, through multimodal profiling of blood and tissues from organ donors from the US and UK spanning six decades of adult life. Our findings uncover the basis for the complexity of the immune system and how immune cell identity is shaped by tissue and age. We reveal how tissue imparts key features of immune cell composition, activation, and function and how the immune system alters with age through localized changes to specific immune subsets along with generalized alterations in immune cell capacities. In this way, the immune system provides a highly resilient and adaptive form of protection optimized for the diverse environments of the body and the numerous sources of tissue damage.

The immune system consists of diverse cell lineages and subsets that are found not only in specialized lymphoid organs and blood, but also in diverse mucosal and barrier sites. We show that tissue site is a dominant driver of the immune landscape, determining immune cell composition, cell states, and functional capacity. Of the sites profiled here, intestines had the most distinct immune compartment both in terms of composition and gene expression profiles even compared to lungs, another mucosal/barrier site. In particular, intestinal lymphocytes were mostly TRM (for T cells) and plasma cells (for B cells), and gut-specific gene and protein expression profiles included specific homing receptors, and functions such as IL-17 production. Similarly, LN also exerted specific functional and differentiation properties on immune cells, maintaining them in more quiescent and stem-like states. By contrast, lungs, spleen, BMA and blood contained lymphocyte populations with more pro-inflammatory profiles, but distinct myeloid lineage cells with tissue-specific properties. Between lineages, T, NK/ILC, and myeloid lineage cells exhibited dominant tissue-specific features while B cells did not, indicating that B cell identity is less influenced by location.

The tissue-specific signatures identified here have important relevance for immune monitoring and immunotherapies. The compartmentalization of subsets and functional responses in tissues compared to blood, suggests that monitoring tissue-directed responses is not readily assessed in circulation. Indeed, in studies of responses to respiratory infection with SARS-CoV-2, the lymphocyte and myeloid cell responses in the respiratory tract were distinct from blood and correlated more with infection outcome ^60,61^. In addition, the striking segregation of gut-specific subsets and profiles indicates that oral administration of vaccines would be optimal for promoting protection to intestinal pathogens, as demonstrated for the success of rotavirus vaccines^62^. Another clinically relevant finding in our tissue data includes the identification of stem-like profiles for T and NK cells in lymph nodes. As expression of *TCF7* and *LEF1* is associated with complete and partial remission in CAR-T immunotherapies ^63^, the isolation of NK and T cells from LN may constitute optimal sources for generating CAR-T for anti-tumor and anti-viral therapies.

Our dataset enabled a novel assessment of how immune cells across the body become altered with age. We found that age-associated changes were for the most part, specific to lineage and site. For macrophages in lungs and lymphoid organs (but not gut), we found a reduction of the *APOE-TREM2* axis with age. TREM2 can have different effects on macrophage function, depending on context; it has been associated with induction of anti-inflammatory M2 function during infection, but can also promote phagocytosis and sustained inflammation in other infectious conditions ^64,65^. Notably, TREM2 expression in microglia has been widely associated with age-driven dementia in Alzheimer’s disease, and more recently, TREM2-macrophages have been associated with immunosuppression and lack of responses to immunotherapy in tumors ^66^. The loss of TREM2 over age may be due to increased binding by ApoE and could serve as an adaptation to restrict macrophage activation or regulation in tissues. This alteration in regulation of macrophage function with age is important to dissect in future studies to determine whether it plays a role in the increased susceptibility to respiratory pathogens and lung cancer with age.

Other age-associated features were specific to lineages in certain sites. T cells and NK cells in circulation were functionally altered with age, expressing higher levels of genes associated with inflammation, cytotoxicity, and migration, reflecting previous findings in blood ^67^. By contrast, T cells resident in tissues (such as the gut and lung) were impaired in their ability to produce pro-inflammatory cytokines such as IL-17, which is associated with reduced barrier immunity ^68^, suggesting compromised *in situ* protection over age. These impaired tissue responses may result in reduced immune surveillance of an organ and predispose for tumor formation with age as observed for lung and colon cancer– diseases of the elderly ^69^. The increased systemic inflammation and cytotoxicity of surveilling T and NK populations can have both deleterious effects causing tissue damage, but can also help clear senescent cells ^70,71^. Our results also reveal a potential switch to more T cell proliferation and tissue repair functions with age in LN, possibly in response to age-associated tissue damage.

The immune system is distinct in being distributed throughout the body, in its ability to establish memory as resident populations in diverse anatomic sites, and in its ability to adapt with each pathogen encounter and environmental exposure ^8^. In order to meet the complex demands of the host for protection from pathogens and mediators of tissue injury, immune cells require plasticity but also stability, to prevent dysregulated responses that can lead to tissue damage. Through our extensive and deep profiling of immune lineages across organs and tissue systems, we show how immune cell identity is layered, with lineage being the foundation upon which function and fate are determined, followed by location playing major roles in subset delineation and function, with more subtle changes occurring with age. We reveal that lymphocytes undergo distinct age-associated changes depending on the site; there is increased inflammation in circulation while immune potency appears to be better preserved in lymphoid organs. Together, our dataset, models, and analyzes can serve as a comprehensive and actionable resource to inform targeted immune modulation by site and age in future treatments for infectious, neoplastic, and inflammatory diseases.

## AUTHOR CONTRIBUTIONS

N.Y., D.L.F., P.A.S., J.J., and S.A.T. designed the study; M.K., R.M., K.M., and K.S.-P. obtained tissues from deceased organ donors; S.B.W., D.B.R., P.A.Sz, D.P.C., S.K.H., L.B.J., L.E., D.C. performed tissue dissociation and prepared single-cell sequencing libraries; M.M., C.E., S.B.W., D.R., P.A.S., P.A.Sz, D.P.C., A.R.M., E.R., V.V.P.A analyzed and interpreted the data; D.L.F., N.Y., P.A.S., J.J. supervised experiments; N.Y. and P.A.S. coordinated data analysis; D.L.F., N.Y., P.A.S., J.J, S.A.T, S.B.W., D.B.R., M.M., P.A.Sz., C.E., A.R.M. wrote and edited the manuscript. N.Y., D.L.F., P.A.S., J.J., S.A.T. provided funding. All authors read, provided input on, and approved the manuscript.

## ACKNOWLEDGEMENTS

We thank the deceased organ donors, donor families, the extended Cambridge Biorepository for Translational Medicine (CBTM) team, and the transplant coordinators at LiveOnNY for access to the tissue samples. This work was supported by a Chan-Zuckerberg Initiative Seed Networks for the Human Cell Atlas grant (CZF2019-002452) to N.Y, D.L.F., P.A.S., J.J., and S.T., along with NIH grants AI106697, AI128949 awarded to D.L.F. and P.A.S and NIHR Cambridge Biomedical Research Centre funding to J.J (BRC-1215-20014). D.P.C. was supported by the Columbia University Graduate Training Program in Microbiology and Immunology (T32AI106711). Flow cytometry analysis was performed in the CCTI Flow Cytometry Core supported by NIH S10RR027050 and S10OD020056 and the Cambridge NIHR BRC Cell Phenotyping Hub (Department of Medicine, University of Cambridge). The views expressed here are those of the author(s) and not necessarily those of the NIHR, Department of Health and Social Care, or National Institutes of Health (NIH).

## Conflict of Interest

In the past three years JJ has consulted for, or been a member of a scientific advisory board for Sanofi, Roche and Enhanc3DGenomics, and S.A.T. has consulted for or been a member of scientific advisory boards at Qiagen, Sanofi, GlaxoSmithKline, and ForeSite Labs. She is a consultant and equity holder for TransitionBio and EnsoCell.

## Abbreviations

BLO: Blood
BMA: Bone Marrow
SPL: Spleen
LN: Lymph Node
LNG: Lung
JEJ: Jejunum
TN: naive T cell
TCM: central memory T cell
TEM: effector-memory T cell
TEMRA: terminal effector cell
TRM: Tissue resident memory T cell
TREG: regulatory T cell

---

**Extended data Figure 1:**
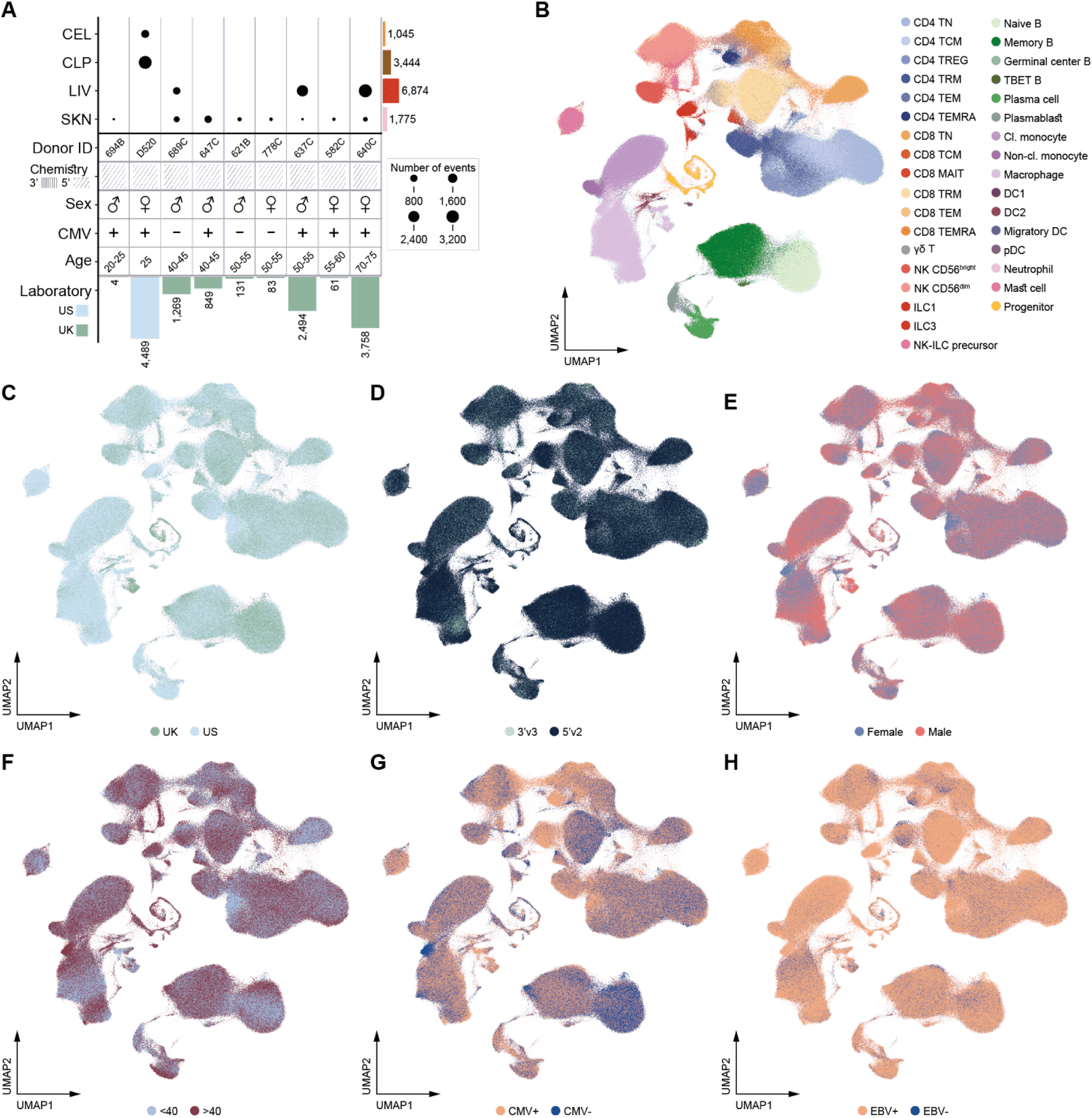
Tissue sites excluded from study and UMAPs of covariates. **(A)** Schematic of four tissue sites collected yet excluded from analysis. **(B)** UMAP of MMoCHi cell annotations. **(C, D, E, F, G, H)** UMAPs colored by collecting laboratory, 10x chemistry, sex, age groups, CMV status, and EBV status.

**Extended data Figure 2:**
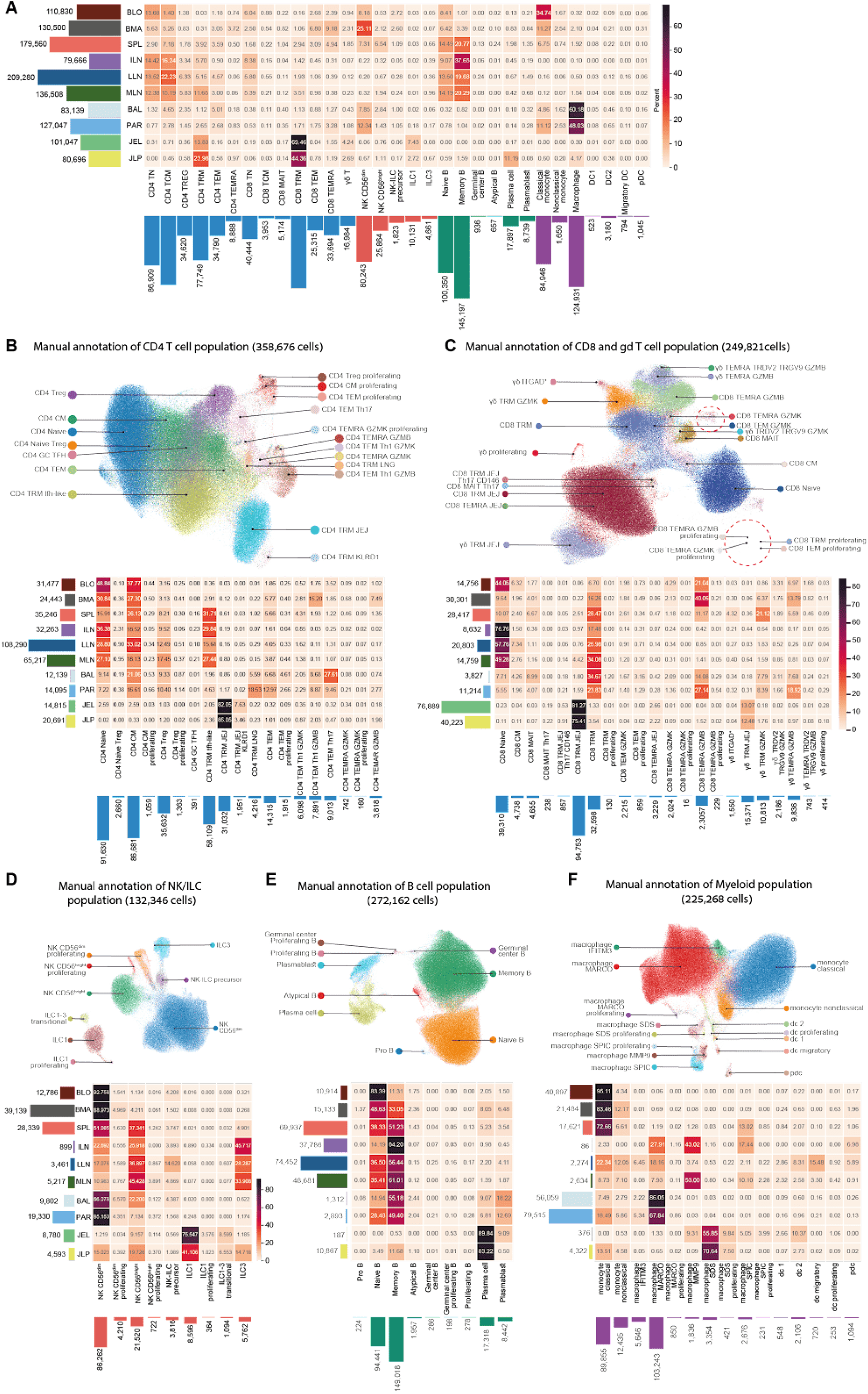
Tissue distribution and refinement of the MMoCHi annotation identifies additional immune populations. **(A)** Heatmap showing the distribution of each MMoCHi classified immune population per tissue (each row sums to 100%), with barplots on the X and Y axis displaying the total number of cells per cell type and tissue, respectively. Manual annotation was performed on each of the MMoCHi classified immune populations and for the **(B)** CD4^+^, **(C)** CD8^+^, **(D)** NK/ILC, **(E)** B cell and **(F)** Myeloid populations the UMAP is colored by annotation label as well as the frequency of each population per tissue within each immune compartment.

**Extended data Figure 3:**
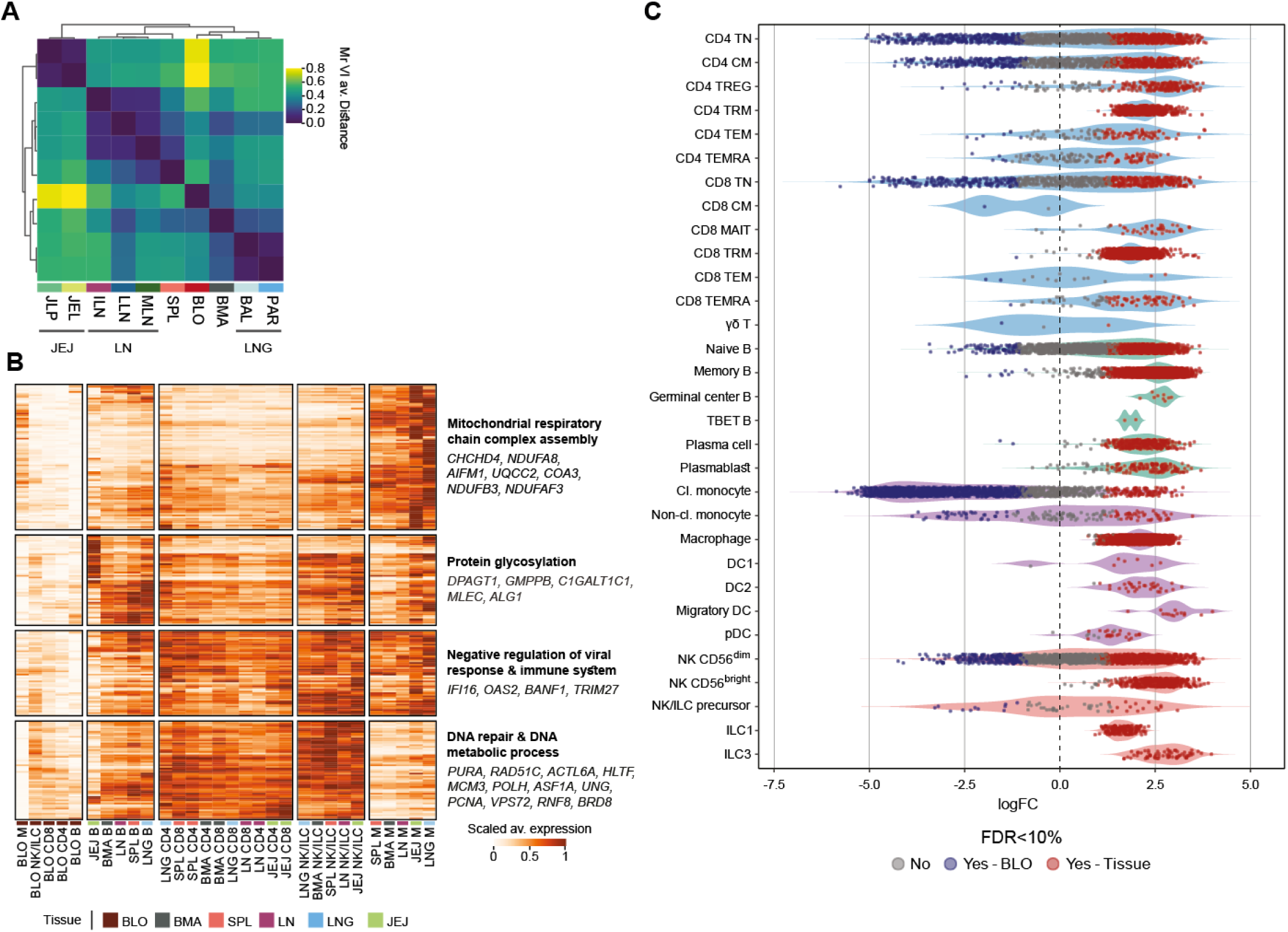
**(A)** Heatmap of distances between tissues. Presented are the average distances, across all immune lineages, as evaluated by MrVI. **(B)** Blood vs. tissue comparative analysis. Analysis was conducted separately for each combination of cell lineage and tissue group. Heatmap displayed normalized expression of genes that were DE (adj. p-value<0.05) with positive LFC in at least 80% of the lineage-tissue combinations. **(C)** Violin plot showing neighborhood differential abundance analysis performed with Milo of blood vs. tissue subsets, which highlights tissue enrichment of specific immune populations.

**Extended data Figure 5.**
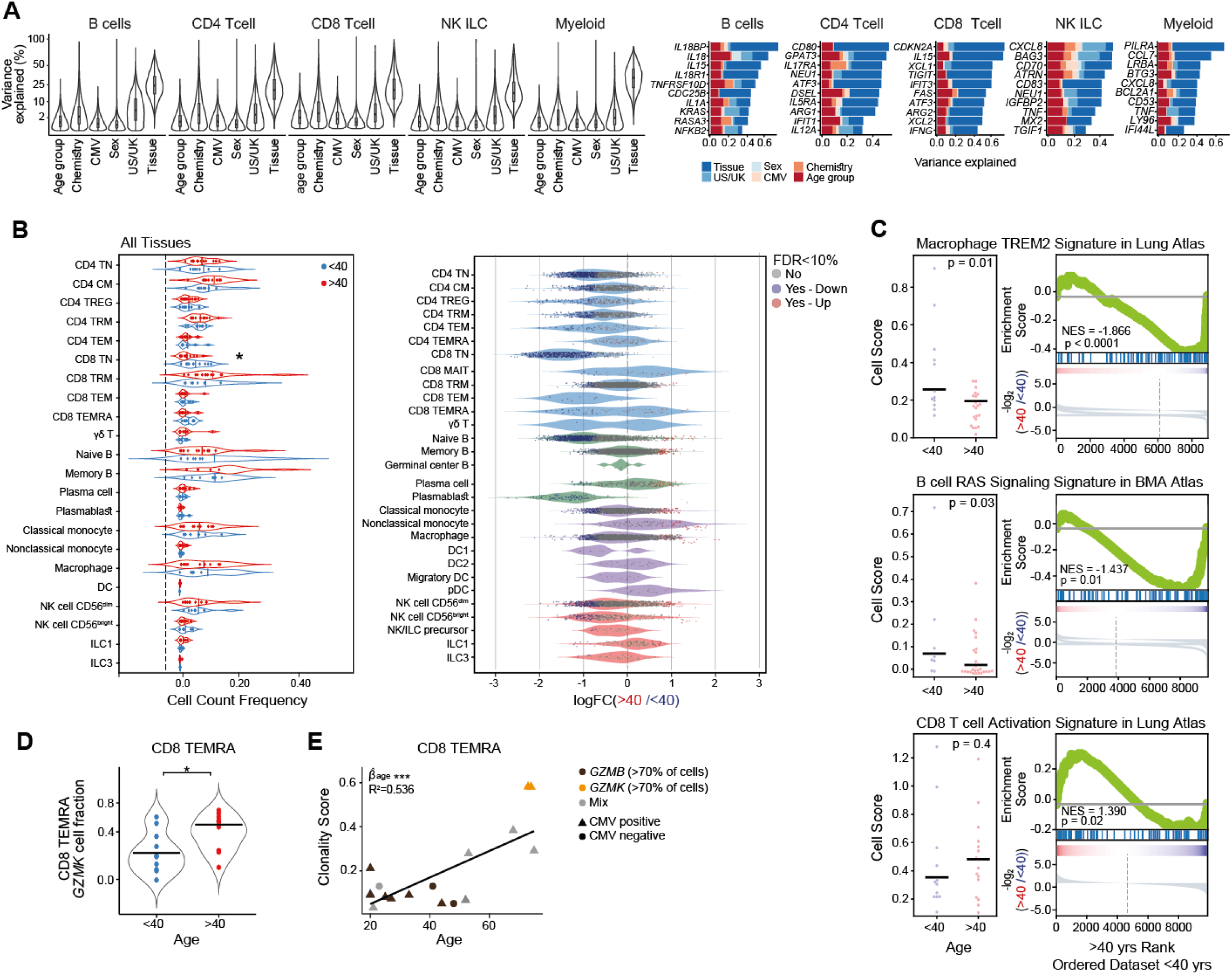
**(A)** Violin plot summarizing variance partitioning analysis, within each immune lineage, separating the fraction of expression variation for each gene into six components including tissue, age group (over vs. under 40 years old), 10x Genomics chemistry (3’ vs. 5’), sex, site of experiment (UK vs. US), and CMV status (positive vs. negative) (left). The variance distribution across these six components in each lineage is displayed for ten representative genes with a high percentage of explained variance by age group (5-20%) (right). **(B)** Changes in cell type abundance with age across all tissues (comparing over vs. under 40 years old donors). (left) Cell type composition is displayed as a violin plot, where each dot represents the frequency of a cell type within each donor (frequencies sum to 100% for each donor). Presented cell types are sufficiently represented (>100 cells) in at least 5 donors in each age group (<40 in blue, >40 in red). Significant differences between age groups are denoted by asterisks (P-values: * <0.05, Wilcoxon test). (right) Violin plot showing neighborhood differential abundance analysis performed with Milo analysis over all tissues. **(C)** scHPF factor cell scores (left) and GSEA analysis (right) showing statistical enrichment of three age-associated factors (top, middle, bottom) among under vs. over 40 year old donors in external datasets. **(D)** Violin plot for the frequency of CD8^+^ TEMRA GZMK^+^ cells out of all CD8^+^ TEMRA cells in blood-rich tissues (BMA, spleen, and blood). Significant differences between age groups (<40 in blue, >40 in red) are denoted in asterisks (P-values: * <0.05, Wilcoxon test). **(E)** Dot plot indicating the clonality score (1−Pielou’s evenness index) of CD8+ TEMRA in blood-rich tissues (each dot represents a donor). The CMV status of the respective donor is indicated by shape, and the fraction of GZMB^+^ and GZMK^+^ in each CD8^+^ TEMRA population is reflected by color. Regression line highlights the positive correlation between age and clonality (R2 = 0.57, p <0.001).

## Notes

### Summary of Updates

The supplementary material has been added

## REFERENCES

1. Szabo, P. A., Miron, M. & Farber, D. L. Location, location, location: Tissue resident memory T cells in mice and humans. Sci Immunol 4, eaas9673. (2019).

2. Terekhova, M. et al. Single-cell atlas of healthy human blood unveils age-related loss of NKG2C+GZMB-CD8+ memory T cells and accumulation of type 2 memory T cells. Immunity 56, 2836–2854.e9 (2023).

3. Tabula Sapiens Consortium* et al. The Tabula Sapiens: A multiple-organ, single-cell transcriptomic atlas of humans. Science 376, eabl4896 (2022).

4. Sikkema, L. et al. An integrated cell atlas of the lung in health and disease. Nat. Med. 29, 1563–1577 (2023).

5. Aizarani, N. et al. A human liver cell atlas reveals heterogeneity and epithelial progenitors. Nature 572, 199–204 (2019).

6. Siletti, K. et al. Transcriptomic diversity of cell types across the adult human brain. Science 382, eadd7046 (2023).

7. Marečková, M. et al. An integrated single-cell reference atlas of the human endometrium. bioRxiv 2023.11.03.564728 (2023) doi:10.1101/2023.11.03.564728.

8. Yosef, N. & Regev, A. Writ large: Genomic dissection of the effect of cellular environment on immune response. Science 354, 64–68 (2016).

9. Thome, J. J. et al. Spatial map of human T cell compartmentalization and maintenance over decades of life. Cell 159, 814–828 (2014).

10. Miron, M. et al. Human Lymph Nodes Maintain TCF-1(hi) Memory T Cells with High Functional Potential and Clonal Diversity throughout Life. J. Immunol. 201, 2132–2140 (2018).

11. Szabo, P. A. et al. Single-cell transcriptomics of human T cells reveals tissue and activation signatures in health and disease. Nat. Commun. 10, 4706 (2019).

12. Poon, M. M. L. et al. Tissue adaptation and clonal segregation of human memory T cells in barrier sites. Nat. Immunol. 24, 309–319 (2023).

13. Dominguez Conde, C., et al. Cross-tissue immune cell analysis reveals tissue-specific features in humans. Science 376, eabl5197 (2022).

14. Kumar, B. V., Connors, T. J. & Farber, D. L. Human T Cell Development, Localization, and Function throughout Life. Immunity In Press, 202–213 (2018).

15. Dogra, P. et al. Tissue Determinants of Human NK Cell Development, Function, and Residence. Cell 180, 749–763 e13 (2020).

16. Kumar, B. V. et al. Human Tissue-Resident Memory T Cells Are Defined by Core Transcriptional and Functional Signatures in Lymphoid and Mucosal Sites. Cell Rep. 20, 2921–2934 (2017).

17. Boyeau, P. et al. Deep generative modeling for quantifying sample-level heterogeneity in single-cell omics. bioRxiv 2022.10.04.510898 (2022) doi:10.1101/2022.10.04.510898.

18. Becht, E. et al. Dimensionality reduction for visualizing single-cell data using UMAP. Nat. Biotechnol. (2018) doi:10.1038/nbt.4314.

19. Caron, D. P., et al. Multimodal hierarchical classification of CITE-seq data delineates immune cell states across lineages and tissues. bioRxiv (2023) doi:10.1101/2023.07.06.547944.

20. Sallusto, F., Lenig, D., Forster, R., Lipp, M. & Lanzavecchia, A. Two subsets of memory T lymphocytes with distinct homing potentials and effector functions [see comments]. Nature 401, 708–712 (1999).

21. Zheng, Y. & Rudensky, A. Y. Foxp3 in control of the regulatory T cell lineage. Nat. Immunol. 8, 457–462 (2007).

22. Thome, J. J. et al. Early-life compartmentalization of human T cell differentiation and regulatory function in mucosal and lymphoid tissues. Nat. Med. 22, 72–77 (2016).

23. Provine, N. M. & Klenerman, P. MAIT Cells in Health and Disease. Annu. Rev. Immunol. 38, 203–228 (2020).

24. Miron, M. et al. Maintenance of the human memory T cell repertoire by subset and tissue site. Genome Med. 13, 100 (2021).

25. Yudanin, N. A. et al. Spatial and Temporal Mapping of Human Innate Lymphoid Cells Reveals Elements of Tissue Specificity. Immunity 50, 505–519 e4 (2019).

26. Muramatsu, M. et al. Class switch recombination and hypermutation require activation-induced cytidine deaminase (AID), a potential RNA editing enzyme. Cell 102, 553–563 (2000).

27. Sutton, H. J. et al. Atypical B cells are part of an alternative lineage of B cells that participates in responses to vaccination and infection in humans. Cell Rep. 34, 108684 (2021).

28. Johnson, J. L. et al. The Transcription Factor T-bet Resolves Memory B Cell Subsets with Distinct Tissue Distributions and Antibody Specificities in Mice and Humans. Immunity 52, 842–855 e6 (2020).

29. Weisel, N. M. et al. Comprehensive analyses of B-cell compartments across the human body reveal novel subsets and a gut-resident memory phenotype. Blood 136, 2774–2785 (2020).

30. Ergen, C. et al. Consensus prediction of cell type labels with popV. bioRxiv 2023.08.18.553912 (2023) doi:10.1101/2023.08.18.553912.

31. Hoffman, G. E., et al. Efficient differential expression analysis of large-scale single cell transcriptomics data using dreamlet. bioRxiv (2023) doi:10.1101/2023.03.17.533005.

32. Mackay, L. K. et al. Hobit and Blimp1 instruct a universal transcriptional program of tissue residency in lymphocytes. Science 352, 459–463 (2016).

33. Milner, J. J. et al. Heterogenous Populations of Tissue-Resident CD8(+) T Cells Are Generated in Response to Infection and Malignancy. Immunity 52, 808–824 e7 (2020).

34. Zaiss, D. M. W., Gause, W. C., Osborne, L. C. & Artis, D. Emerging functions of amphiregulin in orchestrating immunity, inflammation, and tissue repair. Immunity 42, 216–226 (2015).

35. Levitin, H. M., Zhao, W., Bruce, J. N., Canoll, P. & Sims, P. A. Consensus scHPF Identifies Cell Type-Specific Drug Responses in Glioma by Integrating Large-Scale scRNA-seq. bioRxiv (2023) doi:10.1101/2023.12.05.570193.

36. Shan, Q. et al. Tcf1 and Lef1 provide constant supervision to mature CD8+ T cell identity and function by organizing genomic architecture. Nat. Commun. 12, 5863 (2021).

37. Schuettpelz, L. G. et al. Kruppel-like factor 7 overexpression suppresses hematopoietic stem and progenitor cell function. Blood 120, 2981–2989 (2012).

38. Gray, J. I. & Farber, D. L. Tissue-Resident Immune Cells in Humans. Annu. Rev. Immunol. 40, 195–220 (2022).

39. Lavin, Y. et al. Tissue-resident macrophage enhancer landscapes are shaped by the local microenvironment. Cell 159, 1312–1326 (2014).

40. Bohlen, C. J., Friedman, B. A., Dejanovic, B. & Sheng, M. Microglia in Brain Development, Homeostasis, and Neurodegeneration. Annu. Rev. Genet. 53, 263–288 (2019).

41. Natri, H. M. et al. Cell type-specific and disease-associated eQTL in the human lung. bioRxiv (2023) doi:10.1101/2023.03.17.533161.

42. Krasemann, S. et al. The TREM2-APOE Pathway Drives the Transcriptional Phenotype of Dysfunctional Microglia in Neurodegenerative Diseases. Immunity 47, 566–581.e9 (2017).

43. Deczkowska, A., Weiner, A. & Amit, I. The Physiology, Pathology, and Potential Therapeutic Applications of the TREM2 Signaling Pathway. Cell 181, 1207–1217 (2020).

44. Ryan, E. J. et al. Dendritic cell-associated lectin-1: a novel dendritic cell-associated, C-type lectin-like molecule enhances T cell secretion of IL-4. J. Immunol. 169, 5638–5648 (2002).

45. Lee, N. Y. S., Li, M., Ang, K. S. & Chen, J. Establishing a human bone marrow single cell reference atlas to study ageing and diseases. Front. Immunol. 14, 1127879 (2023).

46. Oh-hora, M., Johmura, S., Hashimoto, A., Hikida, M. & Kurosaki, T. Requirement for Ras guanine nucleotide releasing protein 3 in coupling phospholipase C-gamma2 to Ras in B cell receptor signaling. J. Exp. Med. 198, 1841–1851 (2003).

47. Takahashi, Y. et al. Novel role of the Ras cascade in memory B cell response. Immunity 23, 127–138 (2005).

48. Pasello, M., Manara, M. C. & Scotlandi, K. CD99 at the crossroads of physiology and pathology. J. Cell Commun. Signal. 12, 55–68 (2018).

49. Mogilenko, D. A. et al. Comprehensive Profiling of an Aging Immune System Reveals Clonal GZMK(+) CD8(+) T Cells as Conserved Hallmark of Inflammaging. Immunity 54, 99–115 e12 (2021).

50. Xiao, B. et al. Rheb1-Independent Activation of mTORC1 in Mammary Tumors Occurs through Activating Mutations in mTOR. Cell Rep. 31, 107571 (2020).

51. McGuire, P. J. Mitochondrial Dysfunction and the Aging Immune System. Biology 8, (2019).

52. Franceschi, C., Garagnani, P., Parini, P., Giuliani, C. & Santoro, A. Inflammaging: a new immune-metabolic viewpoint for age-related diseases. Nat. Rev. Endocrinol. 14, 576–590 (2018).

53. Persad, S. et al. SEACells infers transcriptional and epigenomic cellular states from single-cell genomics data. Nat. Biotechnol. (2023) doi:10.1038/s41587-023-01716-9.

54. Gaublomme, J. T. et al. Single-Cell Genomics Unveils Critical Regulators of Th17 Cell Pathogenicity. Cell 163, 1400–1412 (2015).

55. Cai, Y. et al. Single-cell immune profiling reveals functional diversity of T cells in tuberculous pleural effusion. J. Exp. Med. 219, (2022).

56. Oja, A. E. et al. Trigger-happy resident memory CD4+ T cells inhabit the human lungs. Mucosal Immunol. 11, 654–667 (2018).

57. Crotty, S. Follicular helper CD4 T cells (TFH). Annu. Rev. Immunol. 29, 621–663 (2011).

58. Collier, D. A. et al. Age-related immune response heterogeneity to SARS-CoV-2 vaccine BNT162b2. Nature (2021) doi:10.1038/s41586-021-03739-1.

59. Goronzy, J. J. & Weyand, C. M. Understanding immunosenescence to improve responses to vaccines. Nat. Immunol. 14, 428–436 (2013).

60. Zhang, W. et al. SARS-CoV-2 infection results in immune responses in the respiratory tract and peripheral blood that suggest mechanisms of disease severity. Nat. Commun. 13, 2774 (2022).

61. Szabo, P. A. et al. Longitudinal profiling of respiratory and systemic immune responses reveals myeloid cell-driven lung inflammation in severe COVID-19. Immunity 54, 797–814 e6 (2021).

62. Folorunso, O. S. & Sebolai, O. M. Overview of the Development, Impacts, and Challenges of Live-Attenuated Oral Rotavirus Vaccines. Vaccines (Basel) 8, (2020).

63. Fraietta, J. A. et al. Determinants of response and resistance to CD19 chimeric antigen receptor (CAR) T cell therapy of chronic lymphocytic leukemia. Nat. Med. 24, 563–571 (2018).

64. Matos, A. de O., Dantas, P. H. D. S., Queiroz, H. A. G. de B., Silva-Sales, M. & Sales-Campos, H. TREM-2: friend or foe in infectious diseases? Crit. Rev. Microbiol. 1–19 (2022).

65. Wu, K. et al. TREM-2 promotes macrophage survival and lung disease after respiratory viral infection. J. Exp. Med. 212, 681–697 (2015).

66. Zhang, H. et al. Immunosuppressive TREM2(+) macrophages are associated with undesirable prognosis and responses to anti-PD-1 immunotherapy in non-small cell lung cancer. Cancer Immunol. Immunother. 71, 2511–2522 (2022).

67. Pereira, B. I. et al. Sestrins induce natural killer function in senescent-like CD8(+) T cells. Nat. Immunol. 21, 684–694 (2020).

68. Merino, K. M., Jazwinski, S. M. & Rout, N. Th17-type immunity and inflammation of aging. Aging 13, 13378–13379 (2021).

69. Foster, A. D., Sivarapatna, A. & Gress, R. E. The aging immune system and its relationship with cancer. Aging health 7, 707–718 (2011).

70. Laphanuwat, P., Gomes, D. C. O. & Akbar, A. N. Senescent T cells: Beneficial and detrimental roles. Immunol. Rev. (2023) doi:10.1111/imr.13206.

71. Hasegawa, T. et al. Cytotoxic CD4+ T cells eliminate senescent cells by targeting cytomegalovirus antigen. Cell 186, 1417–1431.e20 (2023).

