## Supplementary Methods for "Multimodal profiling reveals tissue-directed signatures of human immune cells altered with age"

### Tissue acquisition from organ donors

All work was completed under ethically approved studies. Tissue was obtained from deceased organ donors at the time of organ acquisition for clinical transplantation through an approved protocol and material transfer agreement via LiveOnNY, the organ procurement organization (OPO) for the New York metropolitan area[1,2] and through the Cambridge Biorepository for Translational Medicine (CBTM), REC 15/EE/0152 as previously described[3]. Because of the different amounts of tissues and some distinct samples (i.e. skin, liver, and colon) obtained at each location, the protocols for some of the initial digestions and processing differ as described below.

### Tissue processing

*Columbia University Tissue Processing:* Each tissue was subjected to a tissue-specific protocol to maximize mononuclear cell (MNC) recovery and viability across a diversity of sites[3]. A series of detailed protocols are available on protocols.io[4–10]. Blood samples and bone marrow aspirates shared a protocol where they were diluted 1:4 and layered directly onto Ficoll-paque for density centrifugation (1200 x g for 20 minutes at 20°C) and subsequent mononuclear cell isolation.

Spleen samples were mechanically digested with scissors, and mashed and washed through a 100 µm filter with a solution of PBS containing 5% (v/v) FBS and 2 mM EDTA. The single cell suspension was spun down (400 x g for 10 minutes at 20°C), washed with PBS containing 5% (v/v) FBS and 2 mM EDTA, and layered onto Ficoll-paque for a density centrifugation step (1200 x g for 20 minutes at 20°C) and subsequent mononuclear cell isolation.

Lung and all lymph node samples shared a protocol where they were mechanically digested with scissors, and enzymatically digested on a shaker for 30 minutes at 37°C in IMDM Media (Gibco) containing 1 mg/mL Collagenase D (Millipore Sigma) and 0.1 mg/mL DNase (Worthington). After digestion, the tissue was mashed and washed through a 100 µm filter with a solution of PBS containing 5% (v/v) FBS and 2 mM EDTA. The single cell suspension was spun down (400 x g for 10 minutes at 20°C), washed and resuspended with IMDM Media with 10% (v/v) FBS, and layered onto Ficoll-paque for density centrifugation (1200 x g for 20 minutes at 20°C) and subsequent mononuclear cell isolation.

An airway wash, a form of bronchoalveolar lavage, was performed by washing saline into the airway and collecting it with a 50 mL syringe. This sample was treated with 0.25 U/mL Benzonase (Millipore Sigma) for 30 minutes at 37°C, washed and resuspended with IMDM Media with 10% (v/v) FBS, and layered onto Ficoll-paque for a density centrifugation step (1200 x g for 20 minutes at 20°C) and subsequent mononuclear cell isolation.

Jejunum tissue was processed to separate the epithelial layer (EL) from the lamina propria (LP). The process begins by washing the tissue of intestinal contents or chyme with cold PBS containing 5% (v/v) FBS. The EL was stripped by twice incubating the tissue at 37°C on a shaker for 30 minutes with IMDM Media containing 2mM DTT, 10mM EDTA, and 10% (v/v) FBS. After each strip, the media was removed from the tissue, filtered, and washed through a 100 µm filter with a solution of PBS containing 5% (v/v) FBS and 2 mM EDTA to collect the EL fraction, which was stored on ice until the LP fraction had been collected. In order to collect the cells of the LP, after the second stripping step, the tissue was mechanically digested with scissors and enzymatically digested on a shaker for 30 minutes at 37°C in IMDM Media (Gibco) containing 1 mg/mL Collagenase D (Millipore Sigma) and 0.1 mg/mL DNase (Worthington). After digestion, the tissue was mashed and washed through a 100 µm filter with a solution of

PBS containing 5% (v/v) FBS and 2 mM EDTA. At this step, the single cell suspensions of both the EL and LP fractions were spun down (400 x g for 10 minutes at 20°C), washed and resuspended in IMDM Media with 0.25 U/ml Benzonase (Millipore Sigma). The samples were incubated for 30 minutes at 37°C, washed and resuspended with IMDM Media with 10% (v/v) FBS, and layered onto Ficoll-paque for a density centrifugation step (1200 x g for 20 minutes at 20°C) and subsequent mononuclear cell isolation.

Once isolated, all single cell suspensions were centrifuged (400 x g, 10 minutes at 4°C) and washed twice with PBS containing 5% (v/v) FBS and 2 mM EDTA. Cells were counted using the NC-2000 Cell Counter (Chemometec), and 50 million viable cells from each site were treated with Trustain FcX (BioLegend) and FcR Blocking Reagent (Miltenyi). Cells were subsequently labeled for 30 minutes at 4°C with biotinylated anti-CD66B, anti-CD235ab, anti-CD326 to remove granulocytes, red blood cells, and epithelial cells respectively via streptavidin-coated magnetic particles and negative selection (Bangs Laboratories). Finally, all single cell suspensions were subjected to dead cell removal using a Dead Cell Removal Kit (Miltenyi).

*University of Cambridge Tissue Processing:* Each tissue was subjected to a tissue-specific protocol to generate a single cell suspension of immune cells that has been published at protocols.io[11]. Blood and bone marrow aspirates (sternum) shared a protocol where they were diluted 1:1 with PBS and layered directly onto Ficoll-paque for density centrifugation (1200 x g for 20 minutes at 20ºC) and subsequent MNC isolation.

Spleen and all lymph nodes shared a protocol and were mechanically dissociated by mashing through a 70 µm filter with a solution of X-VIVO-15 (Lonza) containing 1% FBS. The single cell suspension was spun down (600 x g, 10 minutes at 20ºC) and resuspended in X-VIVO-15 containing 1% FBS and layered directly onto Ficoll-paque for density centrifugation (1200 x g for 20 minutes at 20ºC) and subsequent MNC isolation.

Lung and liver shared a protocol and were mechanically digested with scissors into <0.5 cm pieces and transferred to a gentleMACS (Miltenyi) tube containing 2.5 mL X-VIVO-15 and 2.5 mL collagenase. The tube was loaded onto the gentleMACS instrument and a program run that loops three times through a series of short mixing steps (ramp 900 rpm 12 seconds, spin 700 rpm 1 second, ramp 1000 rpm 8 seconds, spin 1500 rpm 1 second, spin 1900 rpm 4 seconds, spin

1500 rpm 1 second, spin 1900 rpm 3 seconds) followed by a longer incubation at 37ºC with gentle mixing (loop twice through spin 50 rpm 15 min, spin 350 rpm 20 seconds) taking 32 minutes in total. We added ~20 µL 0.5 M EDTA to give a final concentration of 2 mM to neutralize the collagenase and passed the digested tissue through a 70 µm filter with a solution of X-VIVO-15 containing 1% FBS. The single cell suspension was spun down (600g x 10 minutes at 20ºC) and resuspended in X-VIVO-15 containing 1% FBS and layered directly onto Ficoll-paque for density centrifugation (1200 x g for 20 minutes at 20ºC) and subsequent MNC isolation.

Jejunum tissue was processed to isolate cells from the epithelial layer (EL) and the lamina propria (LP) and is based on a previously published protocol[12]. The jejunum was first washed in PBS containing 0.04% BSA to remove any chime and then mechanically digested with scissors into pieces <0.5cm. These jejunum pieces were transferred to a 50 mL tube containing 10 mL of X-VIVO-15 containing 2 mM DTT, 5 mM EDTA and 1% FBS and incubated at 37ºC for 20 minutes. This was repeated twice and the chemical digestion solution contains the EL cells and was passed through a 70 µm filter with a solution of X-VIVO-15 containing 1% FBS. The remaining undigested jejunum tissue was transferred to a gentleMACS tube containing 2.5 mL X-VIVO-15 and 2.5 mL collagenase. The tube was loaded onto the gentleMACS instrument and a program run that loops three times through a series of short mixing steps (ramp 900 rpm 12 seconds, spin 700 rpm 1 second, ramp 1000 rpm 8 second, spin 1500 rpm 1 second, spin 1900 rpm 4 seconds, spin 1500 rpm 1 second, spin 1900 rpm 3 seconds) followed by a longer incubation at 37ºC with gentle mixing (loop twice through spin 50 rpm 15 minutes, spin 350 rpm 20 seconds) taking 32 minutes in total. We added ~20 µL 0.5 M EDTA to give a final concentration of 2 mM to neutralize the collagenase and passed the digested tissue through a 70 µm filter with a solution of X-VIVO-15 containing 1% FBS resulting in the LP cells. Both the EL and LP single cell suspension were spun down (600 x g, 10 minutes at 20ºC) and resuspended in X-VIVO-15 containing 1% FBS and layered directly onto Ficoll-paque for density centrifugation (1200 x g for 20 minutes at 20ºC) and subsequent MNC isolation.

Skin was processed using a protocol developed by the Haniffa lab[13]. The subcutis layer was carefully removed from the skin tissue, the remaining skin was digested in dispase for 2-3 hours at 37ºC to allow the epidermal layers to be peeled from the dermal layer. Both the epidermal and dermal layers were washed in PBS to remove any residual dispase and then digested in collagenase at 37ºC overnight. Following collagenase digestion, we added ~20 uL 0.5 M EDTA to give a final concentration of 2 mM to neutralize the collagenase and passed the digested tissue through a 70 μm filter with a solution of X-VIVO-15 containing 1% FBS. The single cell suspension was spun down (600 x g, 10 minutes at 20ºC) and resuspended in X-VIVO-15 containing 1% FBS and layered directly onto Ficoll-paque for density centrifugation (1200 x g, 20 minutes at 20ºC) and subsequent MNC isolation.

### Hashtag and CITE-seq labeling

*Columbia University Protocol:* Each single cell suspension was hashtagged to allow pooling of samples for loading on the 10x Genomics Chromium instrument. Approximately one million MNC per tissue were transferred into 4 mL flow cytometry tubes. Cells were centrifuged at 400 x g for 5 minutes, 4°C, supernatant removed, and resuspended in PBS containing 5% (v/v) FBS and 2 mM EDTA. Cells were treated with Trustain FcX (BioLegend) and FcR Blocking Reagent (Miltenyi) to reduce background labeling and incubated at 4°C for 10 minutes. Each sample was spun at 14,000 x g for 10 minutes, and 1 μL of hashtag was added to each tube. The samples were incubated at 4°C for 30 minutes, and subsequently centrifuged at 400 x g for 5 minutes at 4°C and washed three times with PBS containing 5% (v/v) FBS and 2 mM EDTA. 200,000 cells from each sample were added to a single 4mL flow cytometry tube. This tube was centrifuged at 400 x g for 5 minutes, 4°C. Cells from donors D496 and D503 were re-suspended in the TotalSeq A Universal Cocktail (BioLegend) and cells from all other Columbia donors were re-suspended in the TotalSeq C Universal Cocktail (BioLegend) in PBS containing 5% (v/v) FBS and 2 mM EDTA. Regardless of the panel used, samples were incubated at 4°C for 30 minutes, subsequently centrifuged at 400 x g for 5 minutes at 4°C, and then washed three times with PBS containing 5% (v/v) FBS and 2 mM EDTA before re-suspension in a final volume of 1 mL. **Figure 1** shows the tissues procured from each organ donor.

*University of Cambridge Protocol:* Each single cell suspension was hashtagged to allow pooling of samples for loading on the 10x Genomics Chromium instrument. Approximately 500,000 MNC per tissue were transferred into 1.5mL lo-bind DNA tubes. Cells were centrifuged at 400 x g for 5 minutes, the supernatant removed, and resuspended in 50 μl PBS containing 0.04% BSA.

Cells were treated with 5 µl Trustain FcX (BioLegend) to reduce background labeling and incubated at 4°C for 10 minutes. Each hashtag was spun at 14,000 x g for 10 minutes, and 1µL of hashtag was added to each tube. The samples were incubated at 4°C for 30 minutes, and subsequently centrifuged at 400 x g for 5 minutes, and washed three times with PBS containing 0.04% BSA. Equal numbers of cells from each tissue were pooled and depending on how many tissues were being processed, determined the number of tissues in each pool.

The TotalSeq-C Human Universal Cocktail that targeted 137 proteins (BioLegend) was supplied lyophilised, and was reconstituted by adding 27.5µL of Cell Staining Buffer (BioLegend) and vortex briefly. The CITE-seq cocktail was incubated at room temperature for 5 minutes and then spun at 14,000g for 10 minutes to pellet any antibody aggregates. One vial of CITE-seq reagent was used per donor, and was divided proportionally over the hashtag pools of cells and incubated for at 4 °C for 30 minutes, and subsequently centrifuged at 400 x g for 5 minutes, and washed three times with PBS containing 0.04% BSA. The cells were resuspended in 50 0µl PBS containing 0.04% BSA and passed through a 40 µm Flowmi pipette tip filter to remove any clumps of cells.

For both Columbia University and University of Cambridge protocols, CITE-seq antibody panels can be found in **Supplementary Table 1**.

### Single-cell sequencing

For scRNA-seq experiments, single cells were loaded onto the channels of a Chromium chip (10x Genomics). cDNA synthesis, amplification, and sequencing libraries were generated using either the Single Cell 5′ Reagent (v1 and v2) or 3′ Reagent (v3) Kits. TCRαβ and BCR paired VDJ libraries were prepared from samples made with the 5′ Reagent kit. All libraries were sequenced on either an Illumina HiSeq 4000 or NovaSeq 6000 instrument.

### Alignment and preprocessing of CITE-seq and scTCR/BCR-seq using Cell Ranger

Alignment was performed using Cell Ranger v 6.0.0 from 10x Genomics[14] with the appropriate chemistry option (*fiveprime, SC3Pv2,* or *SC3Pv3*). We added the cell hashing antibody and the protein antibody fastqs to a single call of *cellranger read*. Immune receptors (TCR and BCR) were also aligned using Cell Ranger (*vdj* function). For all downstream analysis of immune receptors, we used Dandelion for pre-processing[15]. However, TCR and BCR alignment results from Cell Ranger were used for quality control and filtering of low quality cells (individual cells with both TCR and BCR detected). In cases, in which a single cell had both TCR and BCR reads, the immune-receptor data were discarded and the cell was annotated as a multiplet. For all alignments, we used reference genome *refdata-gex-GRCh38-2020-A* and immune receptor reference *refdata-cellranger-vdj-GRCh38-alts-ensembl-5.0.0*.

### Quality Control

We merged the immune-receptor and expression data into a single object using *scirpy*[16]. For splitting hashed cells into their respective samples, we used hashsolo with default parameters[17]. Cells that were not assigned to a single sample were removed from downstream analysis. One sample from skin and one sample from thymus were removed entirely due to high levels of hashtag multiplets. We removed the thymus completely from downstream analysis as identifying thymus tissue in older individuals is challenging, and we did not identify thymus-specific cells in our sequenced samples (no thymocytes or *AIRE*-expressing cells).

No filtering was performed at this step except filtering cells with fewer than 50 genes detected, as they are unlikely to be representative of the immune composition in a given tissue. Mitochondrial reads were quantified using a sum over all genes starting with *MT-* and ribosomal genes were quantified using all genes starting with *RPS* and *RPL*. For erythrocyte-related counts all genes starting with *HB* as well as *ALAS2* and *EPOR* were quantified. The last two genes were added to also detect erythrocyte precursors. Cells with more than 20% mitochondrial reads were flagged as potentially low-quality but were kept for downstream analysis. Mitochondrial reads were removed from the gene expression object and from downstream analysis. *MALAT1* was filtered out as a highly expressed gene at this step as it can confound downstream results. To exclude a major effect from contaminating ambient counts we processed the data using decontX[18] and found no evidence of a strong bias in our downstream tasks due to contamination. Two samples (one from liver and one from skin) were removed as they contained high ambient counts. DecontX did not correct the problem of ambient counts in the two samples removed (detected by plasma protein genes like ALB in immune cells). We used a Celltypist model[3] to detect erythrocytes in our data set (https://cog.sanger.ac.uk/celltypist/models/Red_Blood_CZI/v1/Red_Blood_CZI.pkl).For detection of doublets, we used Scrublet with a *sim_doublet_ratio* of 10[19].

We performed an initial integration across all samples derived from a single tissue. For this integration, we used an scVI model[20] with following parameters: 10,000 highly variable genes using the *seurat_v3* option[21] in Scanpy, early stopping enabled and 50 epochs, ten epochs for *n_epochs_kl_warmup*, two layers in encoder and decoder, *nb* gene likelihood and a mini-batch size of 256. We found superior integration using those non-default values.

To perform filtering of low-quality events, we used following quality metrics: the probability of a doublet predicted by Scrublet, the probability of a doublet from HashSolo, the percentage of erythrocyte genes as described above, whether a cell contained both TCR and BCR, whether the Celltypist erythrocyte model predicted a cell to be an erythrocyte as well as cells with a total count below 2,000 UMI, 1,200 unique genes, or 200 protein counts. All scores were added to generate a per cell quality metric. To perform filtering, we argued that cells that group together and have evidence of low quality should be removed from downstream analysis. We first used Louvain clustering on the coordinates from scVI latent space using 15 nearest neighbors to cluster the per tissue integrated data with a resolution of 5.0[22]. Every cluster with a median low quality score (described above) of at least one was removed from downstream analysis. Although low quality events in several tissues were retained with this filtering, their frequency was drastically reduced. We additionally checked for tissue-specific cut-offs to remove additional events and removed clusters with a mean low-quality score of 0.3 from all tissues, except for the lung lymph node and jejunum. In those, we increased the threshold manually to recover more high-quality cells. Using a course cell type annotation based on manual annotation of clusters, we identified some cell types that were consistently filtered even though their quality did not appear to be spuriously low by manual inspection. We retained mast cells and hematopoietic stem cells from all tissues, all macrophages from lymph nodes and spleen, all erythrocytes and platelets from bone marrow, and all monocytes from liver. We afterwards found little evidence for additional low quality events on the tissue level.

Next, we concatenated cells from all tissues and computed again 10,000 highly variable genes using the *seurat_v3* option in Scanpy and used the same parameters as described above but selected this time a mini-batch size of 1,024 to accelerate the training process.

We used this integrated latent space to assign initial cell-types and removed all cell-types that were not labeled as immune cells. Additionally, we removed all cells where manual labeling and automatic labeling using MMoCHi (see below) were inconclusive about coarse cell-type identity (e.g. B cell, myeloid, T cell) These events were of low quality by manual inspection. We additionally defined clusters of low quality cells manually after integrating all cells and removed those post-hoc.

### Immune Cell Subset Classification using MMoCHi

To identify canonical immune cell subsets, we used a recently reported, supervised machine learning algorithm called Multi-modal Classifier Hierarchy (MMoCHi)[23]. MMoCHi achieves high classification accuracy for CITE-seq profiles of immune cell subsets based on extensive benchmarking studies[23]. We employed several key features of MMoCHi (v0.2.1) including its simultaneous use of protein- and RNA-level features for classification, compatibility with a pre-defined reference hierarchy of cell subsets and their established markers, and ability to integrate CITE-seq batches with different levels of signal and background from antibody staining. We first normalized the gene expression (GEX) count matrix using $log(10{,}000C_{g,i} / T_{G,i} + 1)$ where $C_{g,i}$ is the counts for GEX feature $g$ in cell $i$ and $T_{G,i}$ is the total counts for all GEX features in cell $i$. Similarly, we normalized the antibody-derived tag (ADT) count matrix using $log(1{,}000C_{a,i} / T_{A,i} + 1)$ where $C_{a,i}$ is the counts for ADT feature $a$ in cell $i$ and $T_{A,i}$ is the total counts for all ADT features in cell $i$. We then batch-corrected the ADT expression distributions for each ADT feature using landmark registration as in previous methods for flow cytometry[24] and CITE-seq[25]. As described in our earlier report[23], we identified landmarks or peaks in the expression distributions for each ADT feature in a given sample by automatic detection of local maxima following kernel density smoothing using the Scipy functions *signal.find_peaks* and *stats.gaussian_kde*, respectively. Following automatic peak detection, we used the MMoCHi's graphical user interface (GUI) to manually adjust peak identification for some markers. Next, we performed curve registration and warping to align the positive and negative peaks for each ADT feature across experimental batches using the scikit-fda function *preprocessing.registrtion.landmark_elastic_registration_warping*.

MMoCHi trains a random forest classifier for annotation. We provided MMoCHi with a hierarchy of immune cell subsets (nodes) and their canonical surface protein- and RNA-level markers (**Supplementary Figure 1, Supplementary Table 3**), and used the markers to identify high-confidence members (cells) of each node for training. For each marker in the hierarchy, MMoCHi automatically proposes threshold expression levels on the batch-corrected data to identify high-confidence marker-positive and negative cells[23]. MMoCHi also supplies a GUI for manually adjusting these thresholds, which we used for a subset of markers. We establish ADT-level thresholds similarly to gating for flow cytometry, whereas GEX-level thresholds primarily capture cells with any detectable marker expression, due to high transcript drop-out rates in scRNA-seq (**Supplementary Table 4)**. Next, we held-out 20% of high-confidence cells for testing and validation, while the remaining 80% were used for training a random forest classifier using the *ensemble.RandomForestClassifier* function in scikit-learn as described previously[23]. For two of the 24 organ donors and a subset of samples from a third donor, we did not perform CITE-seq and only had scRNA-seq profiles. Thus, these samples were excluded from the MMoCHi classification described here. However, we used a k-nearest neighbors approach to transfer the classifier labels to individual cells profiled from these two organ donors. Specifically, we used the *neighbors.KNeighborsClassifier* function in scikit-learn with *n_neighbors* = 10 to construct a k-nearest neighbors graph in the mrVI embedding of the dataset (see below) and classify the remaining cells. Of the subsets, pDCs were identified using two separate nodes on the hierarchy (**Supplementary Figure 1**), as pDCs shared expression with both B cells and myeloid cells. Once classified, the two subsets were merged into a single population of pDCs. The resulting annotation includes 34 immune cell subsets, as illustrated in **Figure 2**. The MMoCHi annotation was used at two separate levels throughout the manuscript, defined as either one of the 34 fine-grained subsets, or grouped into CD4+ T cells, CD8+ and unconventional T cells (including γδ T cells and CD8+ MAIT cells), B cells, NK cells and ILCs, and myeloid cells (including monocytes, macrophages, cDCs, migratory DCs, and pDCs).

### Differential abundance analysis using Milo

Differences in cell abundances associated with tissue (**Supplementary Figure 6**) or age (**Extended Data Figure 5, Supplementary Figure 7**) were tested using the Python implementation of the Milo framework (https://github.com/emdann/milopy) for differential abundance testing[26]. Briefly, a k-nearest neighbors graph was constructed using similarity in the global mrVI embedding ($k$ = 45) and cells were assigned to neighborhoods (*prop* = 0.02 for all tissue analysis and age analysis for lymphoid tissues; *prop* = 0.015 for the all tissues age analysis; *prop* = 0.1 for the age analysis for all the other tissues/tissue groups).

The cell count in neighborhoods was modeled as a negative-binomial generalized linear model using a log-linear model for the effects of tissue or age on cell counts while accounting for chemistry, site, sex, and CMV status and the total number of cells over all neighborhoods as confounding covariates in differential abundance testing. For age modeling, tissue was also included as covariate for the analysis that included all tissues, but not for analysis of individual tissues. Multiple testing was controlled for using the weighted Benjamini-Hochberg correction, as described in previously[26]. Neighbors with spatial false discovery rate < 0.1 were deemed significant.

The cells belonging to each sample in each neighborhood were then counted and each neighborhood was assigned a cell-type label based on majority voting of the cells belonging to that neighborhood. Mixed neighbors, where the most abundant label was present in less than 75% of the cells within that neighbor, were not assigned a cell-type label. For the tissue UMAPs, the same approach was used but the initial k-nearest neighbors graph was constructed using the mrVI embeddings for each compartment (see below).

### Differential expression and variance decomposition using dreamlet

For the subsequent analysis, our focus was directed towards samples exhibiting substantial representation across experimental sites, cell types, tissues, and donor ages. This resulted in the inclusion of six tissue groups - blood, bone marrow, spleen, gut (jejunum lamina propria and jejunum epithelium), lymph nodes (inguinal, lung, and mesenteric), and lungs (consisting of BAL and lung), encompassing five primary cell types (myeloid, CD4 T cells, CD8 T cells, B cells and ILC/NK cells) as well as 26 cell subtypes, which were characterized with greater annotation detail (**Figure 2**). The applied covariates included 10x Genomics chemistry (3' vs. 5'), sex (male vs. female), laboratory (Cambridge/UK vs. Columbia/NY), and CMV status (positive vs. negative). Donors were categorized into two binary groups (young vs. old using 40-years as a cutoff). Variance decomposition and differential expression (DE) analysis were performed using linear mixed modeling through the dreamlet R package (version 0.99.26 version)[27].

In **Figure 3**, we performed differential expression analysis (dreamlet) of the gene expression data comparing individual major lineages in one tissue group against the remaining tissue groups. We then define a set of DE genes for each tissue group, taking the genes that were detected (adjusted p-value<0.05, fold-change>2) in three or more of the major immune lineages. We then took the union over all tissue groups and hierarchically clustered the resulting set. For the cell type-tissue groups in **Figure 3**, we computed Spearman correlations between cell type-tissue group profiles and then hierarchically clustered using the *cluster.hierarchy.linkage* function in Scipy with Ward's method and Euclidean distance. For clustering genes, we used *cluster.hierarchy.linkage* function in Scipy with Ward's method and Euclidean distance directly on the normalized gene expression values.

Similarly, we performed differential expression (*scanpy.tl.rank_genes_groups*) on the ADT profiles across tissues on log-normalized data. Analysis is conducted separately for each donor, to limit technical effects of staining. Within a tissue comparison, we took proteins that were significant after a Benjamini–Hochberg adjustment for multiple comparisons (adjusted p-value<0.05, fold-change>1.5) in >65% of donors. As above, we included ADTs that were significant in two of the major lineages within any given tissue (a more lenient criterion compared to genes, due to the lower number of proteins in our data). Protein expression data were then hierarchically clustered using the same approach as described above for hierarchical clustering of gene expression data.

In **Figure 4**, for the differential expression analysis (dreamlet), we evaluated three tissue groups (Gut, Respiratory and Lymph nodes), using one vs. all or one vs. another comparisons, while accounting for the covariates age group, sex, CMV status, chemistry, and site.

In **Figure 5**, we conducted a differential expression analysis to quantify transcriptomic changes between old and young donors in each of the six tissue groups mentioned above. We clustered genes that showed significance (adjusted p-value<0.05) in at least two cell types or tissues. Genes identified as significant with expression levels above 0.1, were further characterized within more extensively annotated cell types. Within clusters 4 to 6 the highly expressed genes were also significant (adjusted p-value<0.05) in at least two cell types or tissues. While clusters 1-3 are more tissue or cell type specific.

In **Extended Data Figure 5**, the variance partitioning analysis estimates precision weights by fitting a linear model for each gene in every primary cell type or higher-resolution annotated cell types. The model's covariates encompass tissues, age group, and additional binary groups (CMV, site, chemistry, sex).

### Identification of gene co-expression patterns using consensus scHPF

For identifying cross-tissue and cross-donor gene signatures for each major immune lineage, we constructed probabilistic factor models directly from scRNA-seq count matrices using scHPF. The output of scHPF includes two matrices - an *M* x *K* gene score matrix containing weights for each of *M* genes in each of *k* factors and a *K* x *N* cell score matrix containing weights for each of *N* cells in each of *K* factors. In the original report of scHPF, the algorithm required a user-supplied value of *K*, the number of factors in the model[28]. Here, we use a new, consensus factorization implementation of scHPF where the user specifies a broad range of *K* values from which many scHPF models are generated[29]. The gene score matrices for these models are then clustered to identify *K* recurrent factors, which are combined to seed a final round of training to construct a final consensus model with *K* factors.

We constructed two types of scHPF models: 1) tissue-level models (**Figure 4**) where the number of cells from each of three tissue types was balanced by random sub-sampling (gut - jejunum epithelium/jejunum lamina propria, lung, and lymph node - mesenteric, lung) and 2) donor-level models (**Figure 5**) where the number of cells from each organ donor was balanced. We constructed both types of models for CD4 T cells, CD8 T cells, NK cells, ILCs, B cells, and macrophages. For donor models, where per donor cell numbers can be limiting for some lineages, we only included donors with at least 300 cells for a given lineage. In both cases, the count matrices were randomly downsampled such that the average number of transcripts per cell was the same for each organ donor to avoid coverage bias. scHPF models considered only protein-coding genes (excluding T cell receptor and immunoglobulin cassettes) detected in at least 1% of cells across the final subsampled and downsampled training matrix.

For all consensus scHPF models, we ran scHPF five times for each of 16 values of $K$ (15-30) from which we selected the top three models for each value of $K$ based on convergence criteria for clustering. We applied walktrap clustering to identify recurrent clusters, which we required to form clusters with factors from at least two different models from which we trained the final consensus model[29].

We sought to identify factors from the tissue-level scHPF models of each major immune lineage that were shared across cell types. As described above, we first constructed consensus scHPF models for CD4 T cells, CD8 T cells, macrophages, NK cells, ILCs, and B cells with equal representation of cells from each of three major tissue types (gut, lung, and lymph nodes). From each model, we removed likely nuisance factors containing heat shock protein-encoding genes (common dissociation artifact, >1 gene), ribosomal protein-encoding genes (common coverage artifact, >10 genes), genes from the highly inducible metallothionein cluster (>1 gene), hemoglobin transcripts (red blood cell contamination, >0 genes), and genes in a previously published signature of dissociation-induced cell stress in scRNA-seq (>7 genes)[30] among the 30 top-weighted genes. Next, we computed the average cell score for each factor in each of the three major tissue types, and identified all factors with an average tissue type score that was at least 80% higher in one tissue type than the average of the remaining two. Thus, the resulting set of 53 scHPF factors from across all six lineage-specific models exhibit some degree of tissue specificity. To compare these factors to each other, we computed the Pearson correlation between the gene score vectors for each pair of factors. We then identified factors with a pairwise correlation that was greater than the 95% confidence threshold with at least two other factors, which yielded 31 scHPF factors from across the six major immune lineages. Finally, we performed hierarchical clustering of the Pearson correlation matrix for these 31 factors (*clustermap* function in Seaborn with Euclidean distance) to identify modules containing factors with similar gene signatures that originated from different, cell type-specific scHPF models (**Figure 4**).

As described above, we constructed donor-level scHPF models for each major immune lineage with uniform representation of cells from each donor in order to identify age-associated gene signatures, as shown in **Figure 5**. For each scHPF model, we performed linear mixed effects modeling (LMM) to account for covariates and identify age associations (Figure 5). Each LMM contained six categorical covariates as fixed effects. We encoded age as a binary variable (young vs. old using 40-years as a cutoff) along with 10x Genomics chemistry (3' vs. 5'), sex (male vs. female), experimental site (UK vs. US), and CMV status (positive vs. negative). We also considered three tissue groups (mucosal including BAL, lung, jejunum lamina propria, jejunum epithelium; lymph node including inguinal, lung, and mesenteric lymph nodes; blood-rich including blood, bone marrow, and spleen), which required us to select one category (blood-rich) as a held-out variable. Thus, we have two categorical variables for tissue, which effectively represent mucosal vs. blood-rich and lymph node vs. blood-rich. We encoded donor identity as a random effect. We then used the *MixedLM* function in the Statsmodels module to compute LMM coefficients and p-values for each factor in a given scHPF model using the cell scores as response variables and the *multipletests* function in Statsmodels to compute FDRs (Benjamini-Hochberg method).

To cross-validate age-associated scHPF factors in other datasets, we further analyzed a bone marrow atlas containing 36 age-annotated donors[31] with good B cell representation for a B cell aging factor and a lung atlas containing 29 age-annotated donors[32] with good macrophage and CD8 T cell coverage (**Extended Data Figure 5**). Although CD8 T cells can be detected in many tissues, the CD8 T cell aging signature was particularly associated with TRM, which are highly abundant in mucosal tissues such as lung. Using the published cell type annotations from each atlas, we extracted the appropriate scRNA-seq profiles and projected them into the corresponding donor-level scHPF models generated from the data reported here using the scHPF *project* function. This resulted in cell scores for cells from the external data sets for the same factors that were generated from this data set, allowing us to compare the average cell scores for young vs. older donors from the external data. As an orthogonal approach, we also performed pseudo-bulk differential expression analysis between older and younger donors (using an age cutoff of 40 years) from the external data sets, ranked the genes by fold-change, and used gene set enrichment analysis (GSEA) to analyze the statistical enrichment of age-associated factors among young vs. old donors. We used the top 100 genes (ranked by scHPF gene score) in each age-associated factor as gene sets for GSEA.

### Continuous analysis of aging effects at a single cell resolution (kNNage)

To estimate the changes occurring gradually with age at the single-cell level, we computed the age for each cell as a weighted average of donor age for the *k* nearest cells based on the mrVI *U*-latent space. This gave rise to an age coefficient (kNNage) for each cell, which lies in the range of ages of the donors in our dataset. Here, $k$ was defined as min($N_{tot}^{0.5}$, 100) where $N_{tot}$ is the number of cells and calculated using the *KNeighborsRegressor()* function in scikit-learn. To mitigate the effect of the number of cells per donor in the dataset, the weight for each donor was defined as $N_{tot} / N_d$ where $N_d$ is the number of cells from the donor *d*, and these weights were used to calculate a weighted average for the kNNage for each cell. In subsequent analyses involving kNNage, cells surrounded exclusively by cells from the same donor, with no other cells from a different donor within $k = N_{tot}^{0.25}$ were disregarded (~1.4% of cells). The scAge was calculated separately for each cell group and tissue group.

Correlation between gene expression and kNNage was calculated on scaled (counts per 10,000) and log1p expression values for each coarse cell group and tissue group as well as for fine cell types and tissue group. For each cell type and tissue group, Pearson correlations between kNNage and gene expression were computed considering only approximately 12,000 genes from the differential expression analysis. To evaluate significantly correlated genes, we calculated empirical p-values for the correlation coefficient for each gene. This was done by repeating the kNNage calculations and subsequent Pearson correlations *R* while randomly assigning ages to each donor at the start. In total, there were 259 random shuffles for fine cell groups and 159 shuffles for coarse cell groups, plus the calculations with the correct age. Empirical p-values were defined as: (number of occurrences of $R^2$ < shuffled $R^2$ +1) / (number of shuffled samples + 1). Genes that have empirical p-values below 0.1 and $|R|>0.1$, and are also detected as significant in the pseudo bulk DE (FDR<0.05). Out of the genes selected in **Figure 5a** (300 genes), 100 genes detected as significant in scAge. Significant genes detected in both pseudo bulk DE and in kNNage are displayed in the kNNage analysis in **Figure 5**.

### Integration, cell state embedding, and age-counterfactuals using mrVI

ScVI was not yielding a fully integrated latent space but clustered by site of collection. MrVI uses a mixture-of-gaussian as a prior, which enforces stronger separation of true cell-state and effect of donors on gene expression, as has been recently demonstrated by Boyeau *et al*[33]. mrVI takes advantage of a prior based on a multimodal variational mixture of posteriors (similar to a VampPrior[34]), which have been shown to outperform Guassian priors for scRNA-seq integration in benchmarking studies[34]. Briefly, mrVI finds a sample-agnostic latent space $U$ and computes a sample-specific embedding. A second latent space $Z$ is defined by adding an attention-based concatenation between $U$ and the sample embedding space to the original $U$-space. Another layer of attention is used to incorporate an embedding of 10x Genomics chemistry and experimental site (Cambridge/UK vs. Columbia/NY), and this third latent space is decoded using a linear decoder to yield the rate of a negative binomial distribution. We use a cell type-aware Gaussian mixture prior in $U$-space. To introduce cell type awareness, we use a bias to the mixture proportions that makes it likely for cells of the same type to be sampled from the same Gaussian.

For the latent embedding highlighted throughout the manuscript and used for manual cell-type curation, we used the donor keys as the sample keys and used the output of the cell-type classifier described above as the cell-type prior in mrVI. For the mrVI differential gene expression analysis, we used a string concatenation of donor and tissue as sample key (our goal was to extend the analysis looking at tissue specific effects of the sample embedding) and we used the cell-type groups used for the factor model (described below) for the bias on cell-type information. For both models, we used default parameters except *n_epochs_kl_warmup* of 25, *n_latent_u* of 20, *n_latent* in $Z$-space of 200, dropout in *qz* as well as *pz* of 0.03 (adopted from [33], follow-up manuscript under preparation).

For the differential expression analysis described in **Figure 6**, we subset the sample embeddings to a respective tissue group and modeled the predicted ε by a mixed effect linear model adjusting for covariates in sex, CMV status, and age group (split into two groups above and below 40). The effects of 10x Genomics chemistry and site were corrected with a random effect. A ridge regression parameter of 0.1 due to collinearity of cofactors was added. This decomposition of ε was performed for every single cell. This yields an estimated effect in $Z$-space for each covariate.

The effect vector was added to the mean cell embedding in *Z*-space and differential gene expression was computed based on the modified and mean embedding for each cell. For downstream analysis, this matrix of estimated log-fold-changes for each cell and gene was further processed for each coarse cell-type. First, all cells were filtered out that were represented only in fewer than three samples[33]. Second, genes were filtered to only retain genes with an average expression in that cell-type of above 0.01 and an estimated log-fold-change with a 95-percentile above 0.1 (we want to retain only genes that might be affected by age in a group of cells). To dissect those predicted gene effects into modules, neighborhood smoothing was performed using 15 nearest neighbors in *U*-space and multiplying two times the normalized affinity matrix with the predicted gene effects. Spectral co-clustering was afterwards performed with four gene clusters and four cell clusters using the mini-batch enabled version in scikit-learn[35]. Marker genes for each module were identified by averaging the predicted log-fold-changes across all cells from the corresponding cell module and top 50 genes for each module were identified. We used decoupleR to first compute a module score of log fold change scores using weighted means of the signs of those marker genes[36]. We additionally computed the weighted means (weighted by average log-fold-changes) of the raw expression values.

To confirm our findings on a per-gene level in the automatic reports, we used pseudobulk estimates of differential expression filtering all samples represented in a gene module with fewer than 10 cells and using covariates here as age group, site, and sex and filtering out all genes with a total count below 20. For pseudobulk computation, we used pyDESEQ2[37]. Filtering was loose here and CMV was removed as covariate as cell modules might contain a small number of cells and pseudobulk fails if covariates are collinear.

To report differential abundance in the mrVI report, we compute differential abundance in the mrVI embedding space. In short, we use the mean and variance predicted by the model in *U*-space and compute the probability that the position of a cell is derived from another sample (probability of measuring this event under the aggregate posterior of all cells in another sample). This differential abundance is not corrected for covariates and is meant to give an overview of representation but not for statistical testing. Numbers reported are the log-likelihood difference between all samples from individuals above and below 40 years of age.

In **Figure 6**, we focused our analysis on the gene module inside CD4 T cells that contained TRM. To detect similar cells in other tissues, we computed the best cut-off for the module score to identify cells in a specific cell module based on Youden's J statistic, computed the module score for all cells from other tissues as described above and applied the same cut-off to all other tissues as the tissue we are focusing on. Only the gut contained TRM with a Th17 phenotype and all other tissues had no module-positive cells. We therefore selected all cells with a MrVI predicted positive log-2 fold change of *IL17A* above 0.05. To confirm our findings on a per-gene level, we selected all cells inside a module and used pseudobulk differential expression analysis. We filtered here to all samples with at least five cells and 1,000 counts; we retained genes with total counts of at least three. We corrected for the following covariates: age group, site, tissue, and sex. Manual selection of genes was performed here to form consistent groups of genes that overlap in their function with the mrVI analysis, while genes predicted to be differentially expressed only due to ambient counts were not reported (immunoglobulin genes in CD4 T cells). To report differentially expressed genes in other tissues for a tissue module, a kNN classifier in the *U*-space from **Figure 2** was used to transfer module-positive labels from one tissue to cells from all other tissues and pseudobulk was performed using the same settings in this cell-subset.

### T cell receptor and B cell receptor repertoire analysis using Dandelion

Cell Ranger-mapped TCR and BCR contigs contained in 'all_contigs.fasta' and 'all_contig_annotations.csv' output files were re-annotated using the Dandelion preprocessing pipeline[15]. This pipeline includes the following steps: (1) Sample suffix/prefix assignment to each sample barcode; (2) Re-annotation of contigs with IgBLAST v1.19.0[38] against IMGT (international ImMunoGeneTics) reference sequences (last downloaded: 24/04/2023); (3) Re-annotation of D and J genes separately using blastn to enable the annotation of contigs without the V gene present; (4) Identification and recovery of nonoverlapping individual J gene segments. For BCRs, three additional steps were also performed: (1) Additional re-annotation of heavy-chain constant (C) region calls using blastn (v2.13.0) against curated sequences from CH1 regions of respective isotype class. (2) Heavy chain V gene allele correction using TIgGER v1.0.1[39]; and (3) BCR mutation calling. Cell-level quality control was performed using Dandelion's 'filter_contigs' function, which only considers productive VDJ contigs, asserts that a single cell should only have one VDJ and one VJ pair, or only an orphan VDJ chain, and explicitly removes contigs that fail these checks (except for IgM/IgD and TRB/TRD extra pairs). Contigs that did not match any cell barcodes in the gene expression data were also removed at this step. TCRs and BCRs were then grouped into clones/clonotypes. The following default sequential criteria, which apply to both chain contigs, were applied: (1) identical V and J genes usage; (2) identical junctional CDR3 amino acid length and (3) CDR3 sequence similarity - 100% nucleotide sequence identity at the CDR3 junction for TCRs and 85% amino acid sequence similarity (based on Hamming distance) for BCRs.

TCR or BCR data were then transferred into the corresponding Anndata object. Cells that did not have receptor data or that presented more than one receptor were discarded from further analysis. For T cells, cells previously annotated as MAIT cells or ɣδT cells were also discarded. Clonality of the different populations was calculated as 1−Pielou's evenness index, varying from 0 (more diverse) to 1 (less diverse), with the Pielou's evenness corresponding $H_s$ / $H_{max}$ where $H_s$ is the Shannon entropy of sample *s* and $H_{max} = log_2C$ where *C* is the number of unique clonotypes in *s*. All clonality scores were calculated on a subsample of 100 cells for each donor cell type, tissue, or cell type and tissue. TCR clonotype groups were identified using the Cell2TCR tool (https://github.com/Teichlab/Cell2TCR)[40]. Briefly, TCR motifs were inferred using the *cell2tcr.motifs* function using default parameters. TCR clonotype groups that shared the same motif, contained TCRs from at least two different donors, and were composed of more than 50 cells were selected as examples of groups of TCRs with shared specificity across donors. V(D)J networks for TCR clonotype groups were obtained using dandelion while network plotting and proportion pie charts were obtained using Scirpy[16].

### References

1. Carpenter, D. J. *et al.* Human immunology studies using organ donors: Impact of clinical variations on immune parameters in tissues and circulation. *Am. J. Transplant* **18**, 74–88 (2018).
2. Thome, J. J. C. *et al.* Spatial map of human T cell compartmentalization and maintenance over decades of life. *Cell* **159**, 814–828 (2014).
3. Domínguez Conde, C. *et al.* Cross-tissue immune cell analysis reveals tissue-specific

features in humans. *Science* **376**, eabl5197 (2022).

4. Wells, S. B. Preparation of single cell suspension from human spleen tissue v1. (2021) doi:10.17504/protocols.io.bwq4pdyw.
5. Wells, S. B. Preparation of single cell suspensions of the intra-epithelial layer and lamina propria from human intestinal tissue v1. (2021) doi:10.17504/protocols.io.bwq7pdzn.
6. Wells, S. B. Preparation of a single cell suspension from bronchoalevolar lavage v1. (2021) doi:10.17504/protocols.io.bwrjpd4n.
7. Wells, S. B. Isolation of nucleated cells from bone marrow aspirate v1. (2021) doi:10.17504/protocols.io.bwrupd6w.
8. Wells, S. B. Isolation of nucleated cells from whole blood v1. (2021) doi:10.17504/protocols.io.bwr6pd9e.
9. Wells, S. B. Preparation of single cell suspension from human lung tissue v1. (2021) doi:10.17504/protocols.io.bwr9pd96.
10. Wells, S. B. Preparation of single cell suspension from human lymph node tissue v1. (2021) doi:10.17504/protocols.io.bwsapeae.
11. Rainbow, D., Howlett, S., Jarvis, L. & Jones, J. Multi tissue processing for single cell sequencing of human immune cells v1. (2021) doi:10.17504/protocols.io.bz4qp8vw.
12. James, K. R. *et al.* Distinct microbial and immune niches of the human colon. *Nat. Immunol.* **21**, 343–353 (2020).
13. Reynolds, G. *et al.* Developmental cell programs are co-opted in inflammatory skin disease. *Science* **371**, (2021).
14. Zheng, G. X. Y. *et al.* Massively parallel digital transcriptional profiling of single cells. *Nat. Commun.* **8**, 14049 (2017).

15. Suo, C. *et al.* Dandelion uses the single-cell adaptive immune receptor repertoire to explore lymphocyte developmental origins. *Nat. Biotechnol.* (2023) doi:10.1038/s41587-023-01734-7.

16. Sturm, G. *et al.* Scirpy: a Scanpy extension for analyzing single-cell T-cell receptor-sequencing data. *Bioinformatics* **36**, 4817–4818 (2020).

17. Bernstein, N. J. *et al.* Solo: Doublet Identification in Single-Cell RNA-Seq via Semi-Supervised Deep Learning. *Cell Syst* **11**, 95–101.e5 (2020).

18. Yang, S. *et al.* Decontamination of ambient RNA in single-cell RNA-seq with DecontX. *Genome Biol.* **21**, 57 (2020).

19. Wolock, S. L., Lopez, R. & Klein, A. M. Scrublet: Computational Identification of Cell Doublets in Single-Cell Transcriptomic Data. *Cell Syst* **8**, 281–291.e9 (2019).

20. Lopez, R., Regier, J., Cole, M. B., Jordan, M. I. & Yosef, N. Deep generative modeling for single-cell transcriptomics. *Nat. Methods* **15**, 1053–1058 (2018).

21. Hafemeister, C. & Satija, R. Normalization and variance stabilization of single-cell RNA-seq data using regularized negative binomial regression. *Genome Biol.* **20**, 296 (2019).

22. Blondel, V. D., Guillaume, J.-L., Lambiotte, R. & Lefebvre, E. Fast unfolding of communities in large networks. *J. Stat. Mech.* **2008**, P10008 (2008).

23. Caron, D. P. *et al.* Multimodal hierarchical classification of CITE-seq data delineates immune cell states across lineages and tissues. *bioRxiv* (2023) doi:10.1101/2023.07.06.547944.

24. Hahne, F. *et al.* Per-channel basis normalization methods for flow cytometry data. *Cytometry A* **77**, 121–131 (2010).

25. Zheng, Y., Jun, S.-H., Tian, Y., Florian, M. & Gottardo, R. Robust Normalization and Integration of Single-cell Protein Expression across CITE-seq Datasets. *bioRxiv* (2022) doi:10.1101/2022.04.29.489989.

26. Dann, E., Henderson, N. C., Teichmann, S. A., Morgan, M. D. & Marioni, J. C. Differential abundance testing on single-cell data using k-nearest neighbor graphs. *Nat. Biotechnol.* **40**, 245–253 (2022).

27. Hoffman, G. E. *et al.* Efficient differential expression analysis of large-scale single cell transcriptomics data using dreamlet. *Res Sq* (2023) doi:10.21203/rs.3.rs-2705625/v1.

28. Levitin, H. M. *et al.* De novo gene signature identification from single-cell RNA-seq with hierarchical Poisson factorization. *Mol. Syst. Biol.* **15**, e8557 (2019).

29. Levitin, H. M., Zhao, W., Bruce, J. N., Canoll, P. & Sims, P. A. Consensus scHPF Identifies Cell Type-Specific Drug Responses in Glioma by Integrating Large-Scale scRNA-seq. *bioRxiv* 2023.12.05.570193 (2023) doi:10.1101/2023.12.05.570193.

30. Denisenko, E. *et al.* Systematic assessment of tissue dissociation and storage biases in single-cell and single-nucleus RNA-seq workflows. *Genome Biol.* **21**, 130 (2020).

31. Lee, N. Y. S., Li, M., Ang, K. S. & Chen, J. Establishing a human bone marrow single cell reference atlas to study ageing and diseases. *Front. Immunol.* **14**, 1127879 (2023).

32. Natri, H. M. *et al.* Cell type-specific and disease-associated eQTL in the human lung. *bioRxiv* (2023) doi:10.1101/2023.03.17.533161.

33. Boyeau, P. *et al.* Deep generative modeling for quantifying sample-level heterogeneity in single-cell omics. *bioRxiv* 2022.10.04.510898 (2022) doi:10.1101/2022.10.04.510898.

34. Hrovatin, K. *et al.* Integrating single-cell RNA-seq datasets with substantial batch effects. *bioRxiv* (2023) doi:10.1101/2023.11.03.565463.

35. Dhillon, I. S. Co-clustering documents and words using bipartite spectral graph partitioning. in *Proceedings of the seventh ACM SIGKDD international conference on Knowledge discovery and data mining* 269–274 (Association for Computing Machinery, 2001).
36. Badia-I-Mompel, P. *et al.* decoupleR: ensemble of computational methods to infer biological activities from omics data. *Bioinform Adv* **2**, vbac016 (2022).
37. Muzellec, B., Teleńczuk, M., Cabeli, V. & Andreux, M. PyDESeq2: a python package for bulk RNA-seq differential expression analysis. *bioRxiv* (2022) doi:10.1101/2022.12.14.520412.
38. Ye, J., Ma, N., Madden, T. L. & Ostell, J. M. IgBLAST: an immunoglobulin variable domain sequence analysis tool. *Nucleic Acids Res.* **41**, W34–40 (2013).
39. Gadala-Maria, D., Yaari, G., Uduman, M. & Kleinstein, S. H. Automated analysis of high-throughput B-cell sequencing data reveals a high frequency of novel immunoglobulin V gene segment alleles. *Proc. Natl. Acad. Sci. U. S. A.* **112**, E862–70 (2015).
40. Lindeboom, R. G. H. *et al.* Human SARS-CoV-2 challenge resolves local and systemic response dynamics. *bioRxiv* (2023) doi:10.1101/2023.04.13.23288227.
