## Supplementary figures and images for "Multimodal profiling reveals tissue-directed signatures of human immune cells altered with age"

### Supplementary Fig1

#### Supplementrary Figure 1

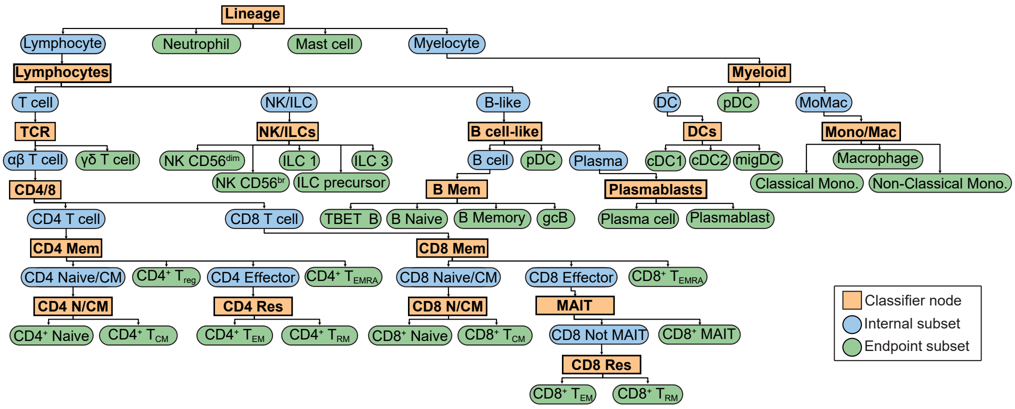

### Supplementary Fig2

**Supplementary figure 2:** Genes and proteins used to define manual annotation

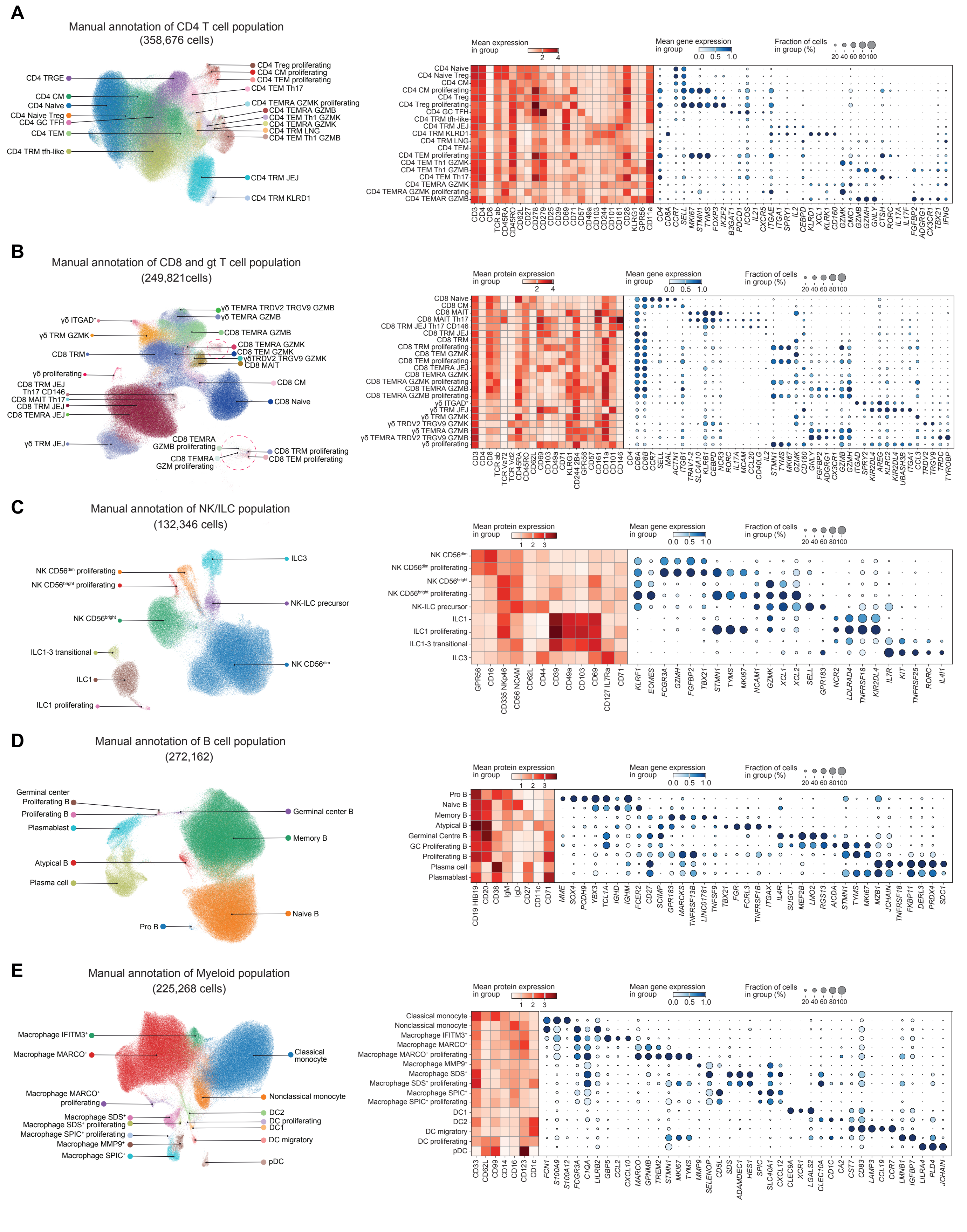

### Supplementary Fig3

**Supplementrary Figure 3**

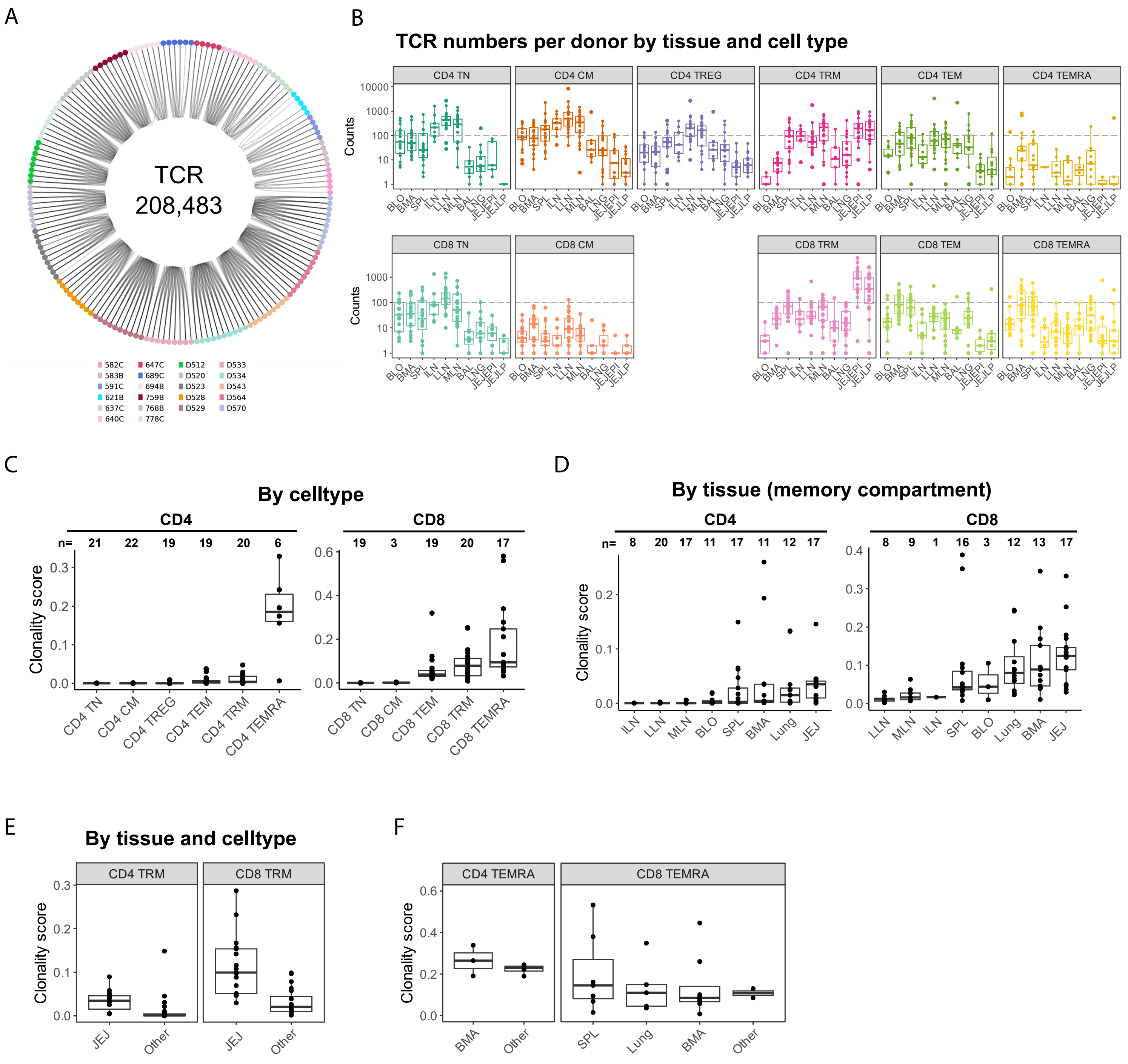

### Supplementary Fig4

#### Supplementrary Figure 4

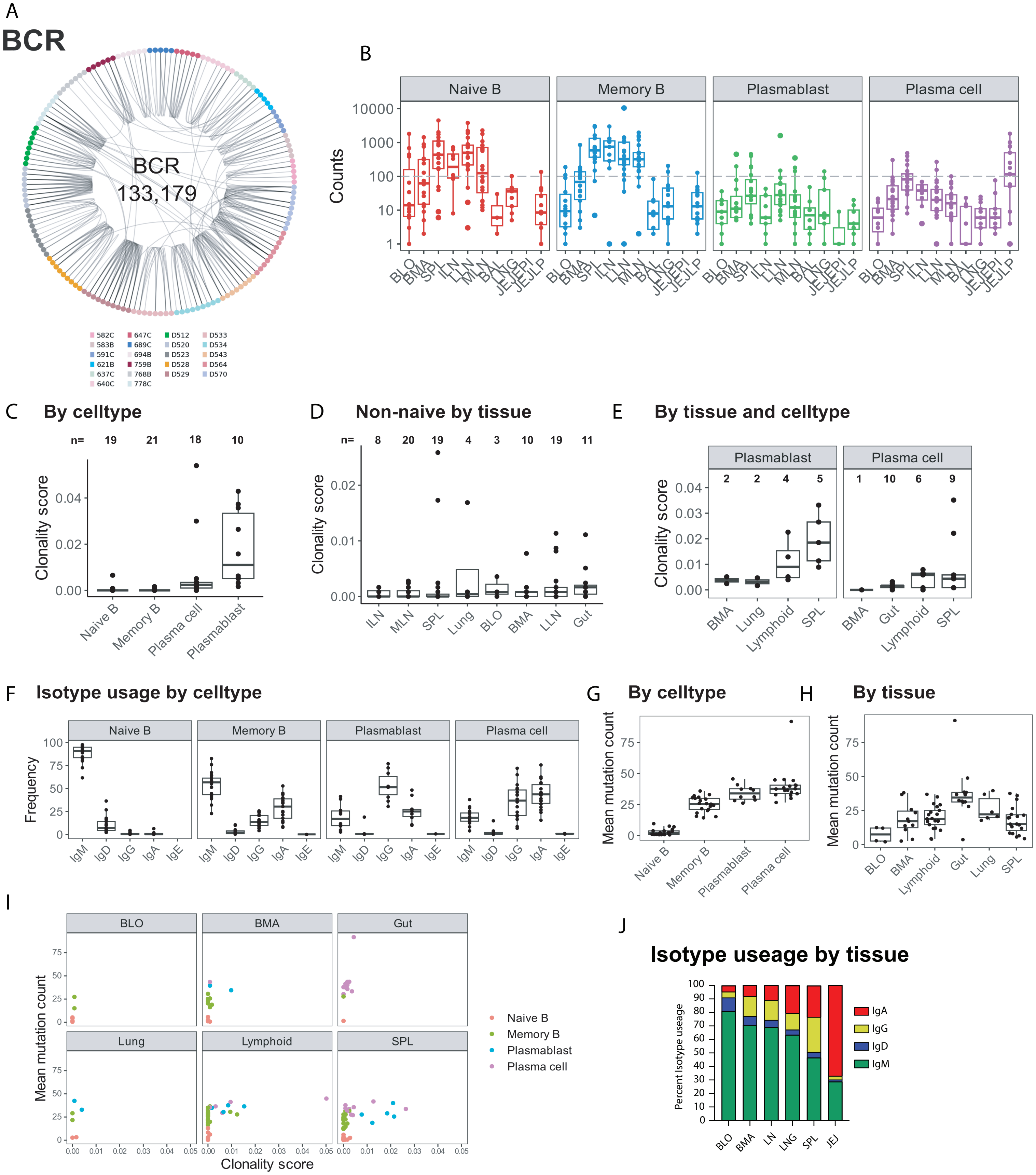

### Supplementary Fig5

##### Supplementrary Figure 5

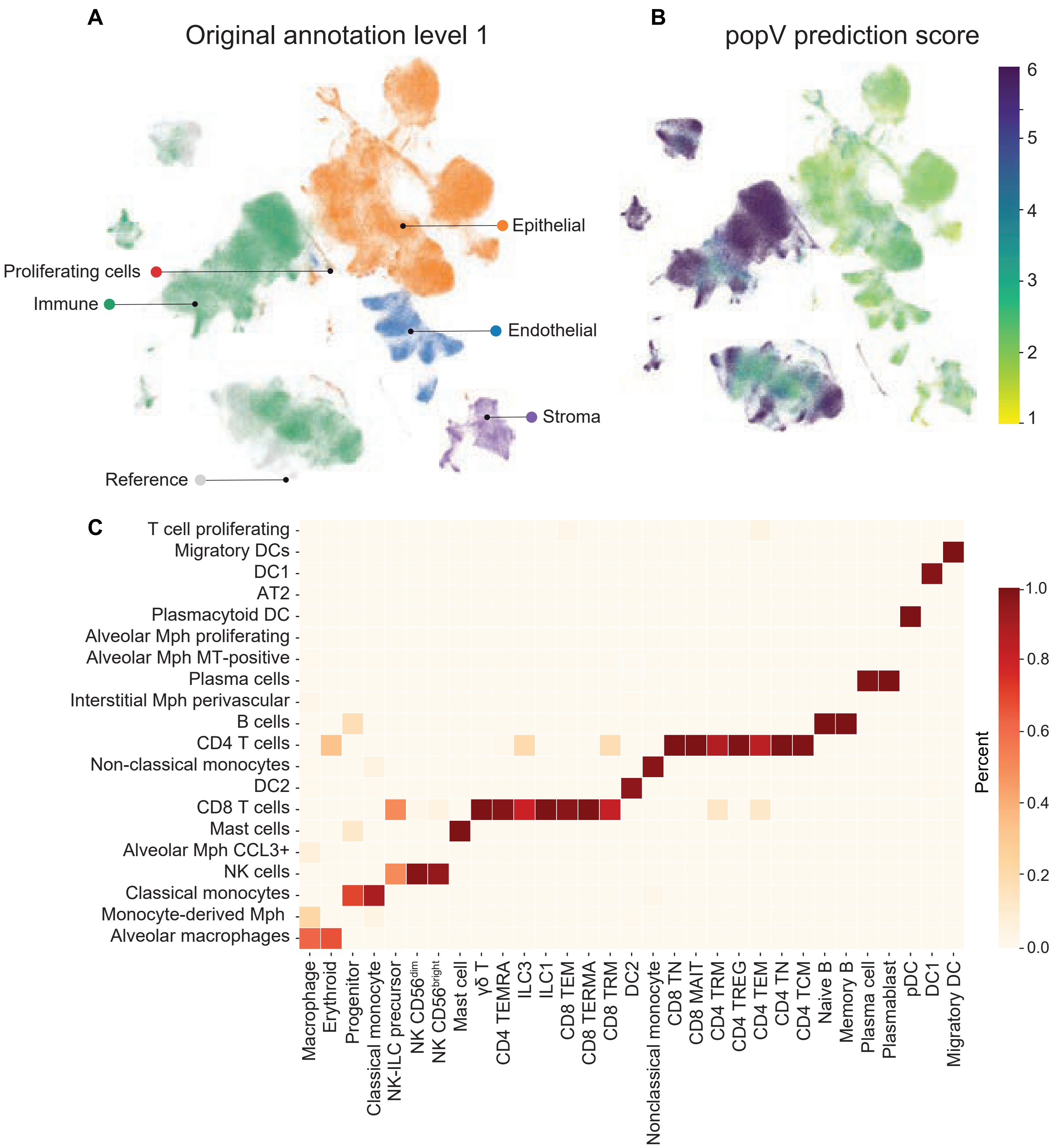

### Supplementary Fig6

#### Supplementrary Figure 6

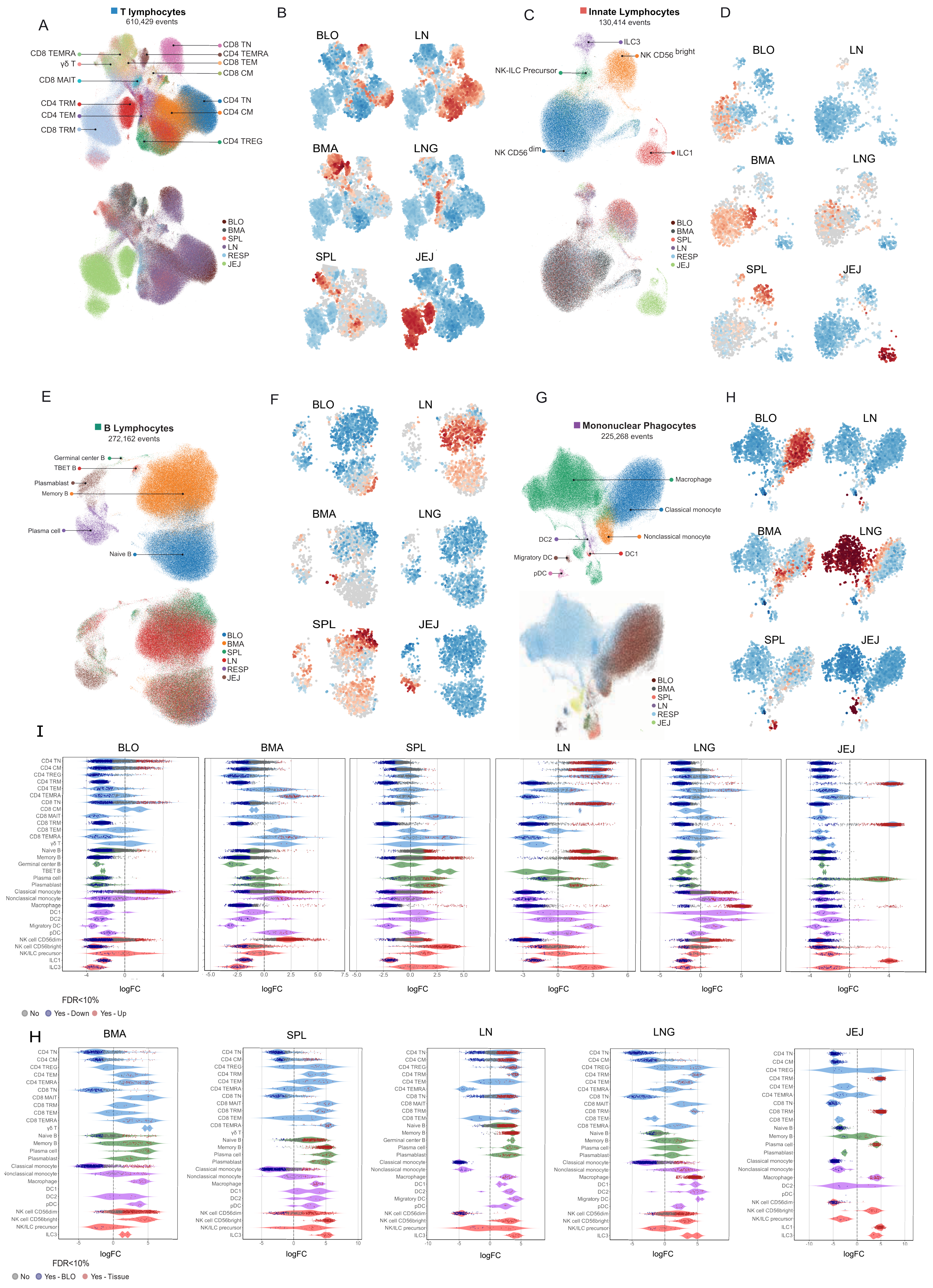

### Supplementary Fig7

#### Supplementrary Figure 7

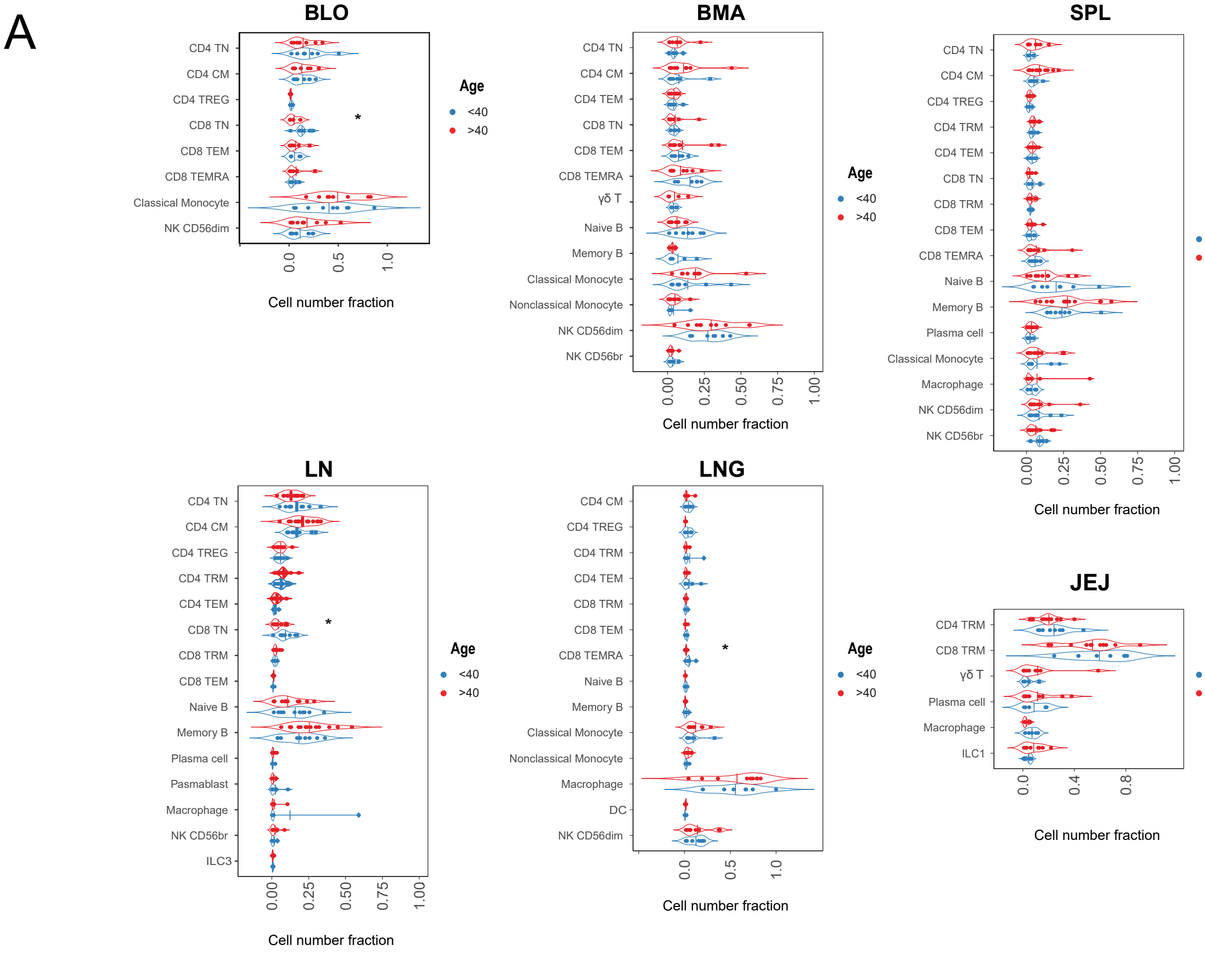


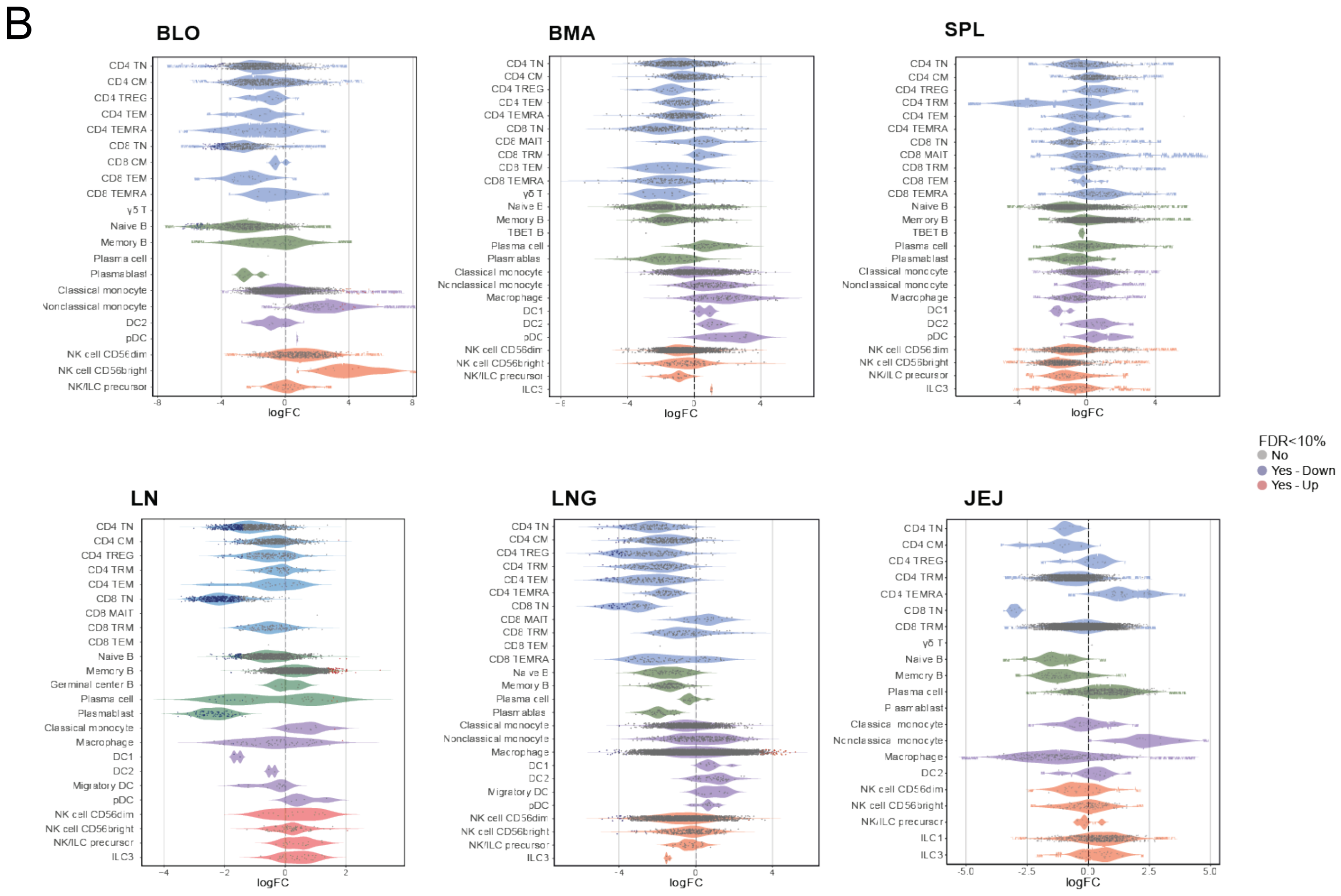

### Supplementary Fig8

#### Supplementrary Figure 8

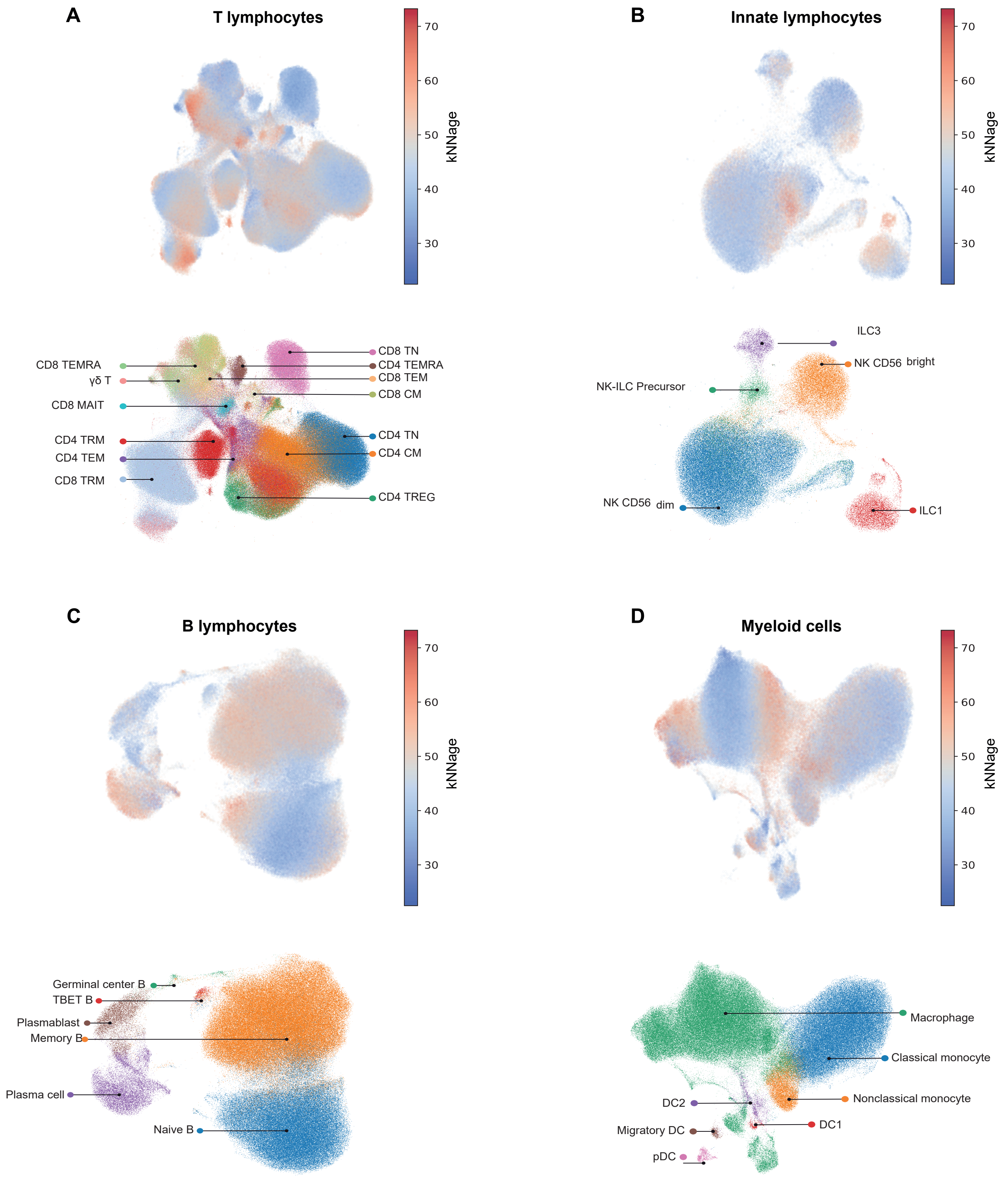

### Supplementary Fig9

##### Supplementrary Figure 9

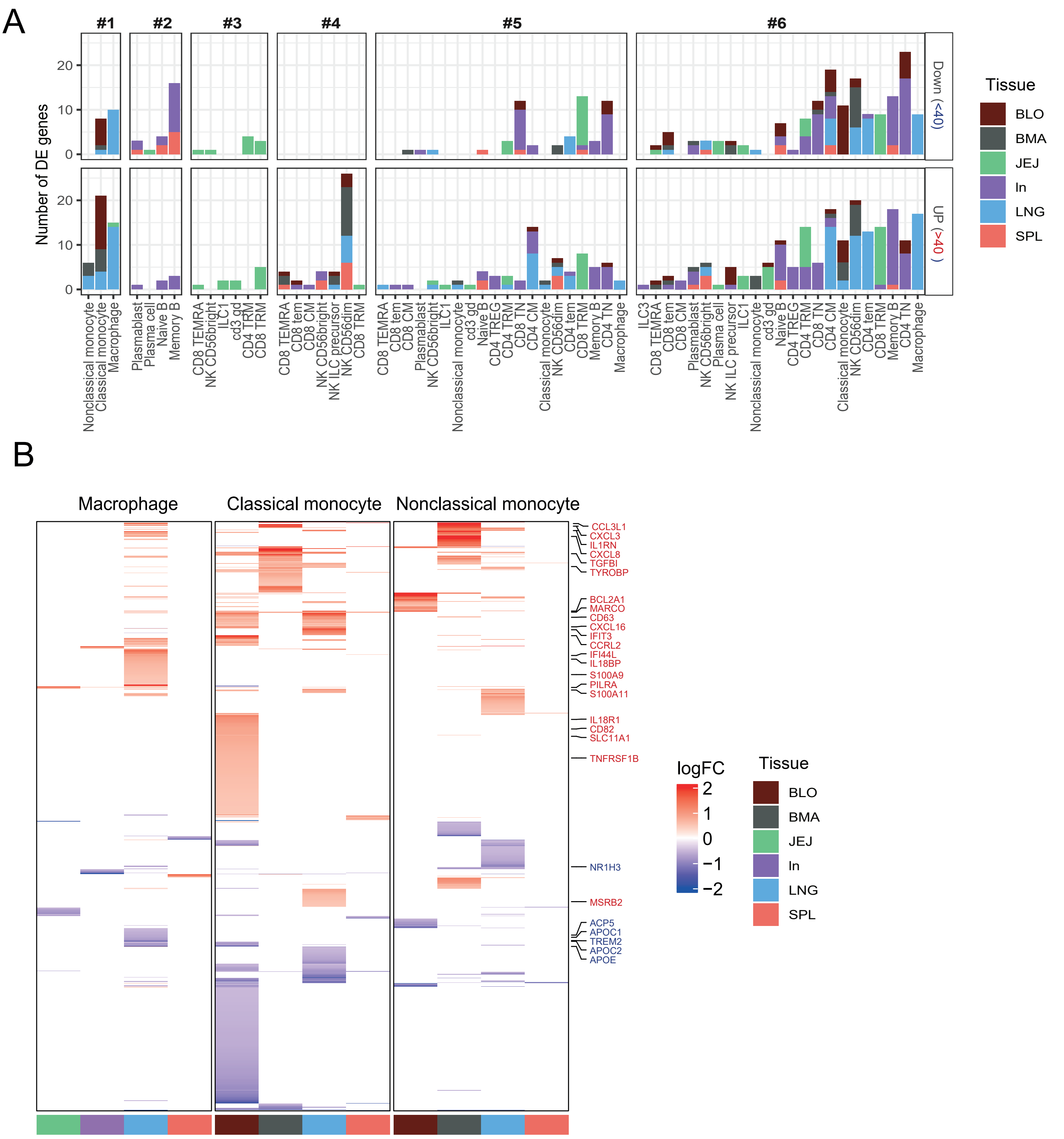
