## Supplementary_Figures_legends for "Multimodal profiling reveals tissue-directed signatures of human immune cells altered with age"

### Supplementary Figure legends

**Supplementary Figure 1 : MMoCHi classifier hierarchy.**

Summary of the cell states considered by our automated annotation process with Multi-modal Classifier Hierarchy (MMoCHi; see Supplementary methods for detail). Labels of cell subsets are organized in a hierarchy. MMoCHi classifiers are applied in each decision point (orange nodes), in the hierarchical order in which they appear, to reach an “end point” annotation (indicated by green nodes).

**Supplementary Figure 2: The genes and proteins used to define the immune populations identified by manual annotation.**

Each immune compartment was manually inspected and additional immune cell populations and states were annotated through sub-clustering. The proteins and genes used to define immune populations in **(A)** $CD4^{+}$ T cells, **(B)** $CD8^{+}$ T cells, **(C)** NK/ILC, **(D)** B cells and **(E)** Myeloid cells are displayed as a red matrix plot (protein) and blue dotplot (genes).

**Supplementary Figure 3: T cell clonality is cell type and tissue specific.**

Summary of the TCR data, based on TCR sequences obtained from 208,483 T cells. **(A)** Circle map showing an overview of the TCR clone sharing information within and across donors. Each dot represents one tissue and is colored by the donor of origin. The vast majority of TCR clones are shared within a donor. **(B)** The boxplots show the number of TCR clones identified from each MMoCHi defined cell population and tissue, with each dot representing a donor. The dotted horizontal line indicated 100 cells which is the minimum cut off for inclusion in the analysis shown on subsequent panels. **(C)** Comparison of clonality scores (1−Pielou’s evenness index) in cell types. Within both the CD4 and CD8 compartment, the TEMRA population has the highest clonality score and the naive population has the lowest clonality score and most diverse repertoire. **(D)** Comparison of clonality scores of the memory cells in tissues. Memory cells in lymph nodes have the lowest clonality score compared to non-lymphoid tissue and bone marrow. **(E)** Comparison of clonality scores of TRMs between gut and other tissues. To account for cellular composition of the tissue, the TCR clonality scores of gut TRM were compared to TRM from all other tissues combined. In both $CD4^{+}$ and $CD8^{+}$ T cell lineages, gut TRMs have a higher clonality score, suggesting a more restrictive TCR repertoire. **(F)** Comparison of clonality scores of TEMRAs across tissues revealed similar levels.

**Supplementary Figure 4: Somatic hypermutations and clonal expansion in plasma and plasmablasts.**

**(A)** Circle map showing an overview of the BCR clone sharing within and across donors. Each dot represents one tissue and is colored by the donor of origin. The BCR sequence data was obtained from 133,179 B cells. The vast majority of BCR clones are shared within a donor. **(B)** The boxplots show the number of BCR cones identified from each MMoCHi defined cell population and tissue, with each dot representing a donor. The dotted horizontal line indicates 100 cells, which is the minimum cut off for inclusion in the analysis shown on subsequent panels. **(C)** Comparison of BCR clonality scores in cell types (numbers of samples appear above each box). The plasmablast population has the highest clonality score. The naive, memory and plasma cell populations have the lowest clonality score and most diverse repertoire. The BCR clonality score was an order of magnitude lower than that of the TCR clonality score. **(D)** Comparison of clonality scores across tissues. To assess how BCR clonality varies across tissue, only non-naive B cells (memory B cells, plasma cell and plasmablast) were used. A similar clonality score was seen across all tissues. **(E)** Comparison of clonality score of plasmablasts and plasma cells in tissues. To account for cellular composition of the tissue, the BCR clonality scores of plasmablast and plasma cell were calculated on tissues with sufficient numbers. In plasmablasts, lymph node and spleen have a higher clonality score compared to lung and bone marrow. A similar trend was seen in plasma cells. **(F)** Comparison of isotype usages in each cell type. From the BCR sequence, we determined the isotypes (IgM, IgD, IgG, IgA and IgE) and showed the frequencies per celltype. Naive B cells are IgM positive, and memory B cells show a proportion being class switched to IgG and IgA. Plasma cells and plasmablast show the highest proportion of IgG and IgA. Comparison of mean somatic mutation counts per cell type **(G)** and per tissue **(H)**. Somatic hypermutations occur in B cell development, and the mean mutation count is lowest in Naive B cells, and increases in B cell maturation resulting with plasma cells and plasmablasts having the highest mutational burden. When the mutation count is plotted per tissue, the gut shows the highest mutational burden, but also has the highest proportion of plasmablasts. **(I)** Relationship between clonality score and mean mutation count. To determine if there is a relationship between clonality score and mutational count, the two parameters were plotted. Within the spleen and lymph nodes, some of the BCR clones with the highest mutational burden also had the highest clonality score. **(J)** B cell isotype usage per tissue. The proportion of IgA, IgG, IgD and IgM isotype use per tissue was calculated (IgE was excluded as the frequency was less than 0.5% in each tissue). A clear enrichment of IgA usage was observed (~70% IgA+) compared to all other tissues with less than 25% IgA usage,

**Supplementary Figure 5: Leveraging our annotation to re-annotate the Human Lung Cell Atlas.** popV is a tool for cell-type label transfer (from a reference atlas to a query dataset) that employs several annotation algorithms and uses consensus voting to determine annotations and evaluate their confidence. It also calculates joint embeddings of the query and reference datasets, which can be used for visualization of the query data and for other analysis tasks. Our dataset is uniquely suited to serve as an immune-cell reference for popV, owing to the breadth of tissues that are included, the number of human subjects, and the level of annotation. **(A)** We trained popV using our data (using MMoCHiannotations as in **Fig. 2**) as the reference data set and the Human Lung Cell Atlas (HLCA) as the query dataset. Joint scVI embedding (calculated as part of the popV pipeline) is presented by UMAP with the reference cells in grey and the query cells colored by their classification into major cell types (as provided by HLCA; note that these original annotations were not used in our analysis).

**(B)** Cells are colored by popV confidence levels (1: low confidence; 6: high). popV highlights non-immune cells as annotated at low confidence, which is expected as these cells are not present in our dataset. **(C)** We then compared the outcome of our immune cell annotations (columns) to the original HLCA annotations, conducted by the authors of that study (rows). We observed an overall high level of concordance. Specifically, our dendritic cell subset annotations (automated) overlap with the curated HLCA labels, highlighting the quality of label transfer. In other lineages, popV is able to resolve additional subtypes for B cells, NK cells, and T cells. Notably, in labelling T cells (bottom left cluster on the UMAP), popV has relatively low confidence (compared to other immune lineages) since distinguishing between some subtypes (e.g. TCM and naive) is difficult when only gene expression (as available at HLCA) is used.

**Supplementary Figure 6: Differential abundance analysis between tissue sites.** Milo analysis to evaluate the effect of tissue on cell type composition confirms the enrichment of specific immune populations in certain tissues (using multivariate model; see Supplementary Methods). **(A, C, E, G)** The immune classification from Figure 2, is shown along with a UMAP per immune compartment colored by tissue of origin. In parts **(B, D, F, H)** the Milo results are displayed as a neighborhood map, with the red color indicating enriched regions in each tissue. For instance, for the T lymphocytes compartment, we showed each immune classification and tissue in the UMAP space (**A**), and the enriched neighbourhoods in each tissue (**B**). This clearly identifies regions within the UMAP that show tissue enrichment, with the gut in T cells and lung in myeloid cells being the most obvious examples. More subtle regions of tissue enrichment can be seen for example TRMs from the lung. (**I**) Violin plots of the milo results - showing the distribution of effects (LFC) of each tissue and for every subset. **(H)** Differential abundance analysis, separately comparing each tissue to blood (shown are only subsets of cells that are present in the blood or in the compared tissue).

**Supplementary Figure 7: Age-dependent variation in cell type abundance in each tissue.** Analysis of immune cell composition, comparing two age groups (over 40 vs. under 40 years old donors) measured in MMoCHicell population per tissue. **(A)** Violin plots displaying the proportion of each cell subset in each tissue. Each dot represents the frequency of a subset within each donor (frequencies sum to 100% for each donor). Presented are combination of tissue and cell subsets that are present (>100 cells) in at least five donors in each age group (<40 in blue, >40 in red). Significant differences between age groups are denoted by asterisks (P-values: * <0.05, Wilcoxon test).**(B)** Violin plot showing a similar analysis of differential abundance, performed with Milo.

**Supplementary Figure 8: Cell-wise Age coefficients estimated by kNNage.**
UMAPs of T cells **(A)**, NK and ILC cells **(B)** ,B cell **(C)** , and myeloid cell **(D)**, colored by estimated kNNage and by their respective MMoCHicell subset.

**Supplementary Figure 9: DE Genes Detected in Both Cell Lineage States and MMoCHi Cell Populations, Reveal Intrinsic Age-Dependent Change.**

**(A)** Bar plots illustrate the count of significant age depended DE genes identified in core cell types and their corresponding MMoCHi cell populations population, per tissue and hierarchical cluster (see Fig 5A,Clustres: #1 to #6). Genes downregulated with age are presented in the upper panel, while those upregulated are shown in the lower pane. **(B)** Heatmap of significant age-driven differentially expressed genes (adj. p-value < 0.05, |LFC| > 0.1) per tissue in Macrophages, Classical, and Non-Classical Monocytes.

### Supplementary Table legends

**Supplementary Table 1: Cite-Seq panels used to measure surface protein expression.**

Two Cite-seq panels of antibodies were used in this project. The two donors sequenced using 10X Genomics 3' Chemistry had Cite-Seq measured using a 277 protein panel. The donors processed using 10X Genomics 5' Chemistry had Cite-Seq measured using a 137 protein panel available from BioLegend (TotalSeq™-C Human Universal Cocktail, V1.0).

**Supplementary Table 2: Donor metadata.**

Table indicating donors ID, Experimental sites (US and UK), the lung donation status (DCD/DBD), Sex, ethnicity, cause of death, EBV and CMV status.

**Supplementary Table 3: MMoCHi classifier markers.**

Table indicating the cell type ("Subset") hierarchical cell lineage ("Parent"), positive and negative markers (Positive Features/Negative Features). Features derived from gene expression are suffixed with "_gex".

**Supplementary Table 4: MMoCHi thresholds.**

Table indicating the negative and positive MMoCHi thresholds for each protein and gene marker. Features derived from gene expression are suffixed with "_gex".

**Supplementary Table 5: Differential gene expression between tissue sites.**

Pertaining to Fig. 3A. Differentially expressed genes ("ID") in cell lineage ("assay", m= myeloid, cd4,cd8, b, ILC_NK) comparing single tissue to all other tissues ("tissue_groups").

**Supplementary Table 6: Differential protein expression genes for Figure 3B tissue analysis.**

Differentially expressed proteins presented in a similar format as Table 5. Pertaining to Fig. 3B.

**Supplementary Table 7: Gene scores in scHPF factors used for studying variation between tissues.**

Table indicating the genes scores over each tissue factor

**Supplementary Table 8: Differential gene expression between tissue sites, in finer-grained (MMoCHi labels) immune subsets that were highlighted by scHPF.**
Pertaining to Fig. 4D,G,J. MMoCHi cell types differentially expressed genes over tissue groups using pseudo-bulk (Dreamlet); LNG vs JEJ, LN vs mucosal (JEJ+LNG), and JEJ vs LNG and LN.

**Supplementary Table 9: Differential expression analysis to study the effect of age in each immune lineage**.
Differential expression tests, using discrete pseudo-bulk analysis with Dreamlet (comparing >40 vs. <40 year old donors), and continuous single-cell analysis with kNNage. Analysis is done separately for every immune lineage.

**Supplementary Table 10: Differential expression analysis to study the effect of age in each finer-grained (MMoCHi labels) cell subset**.
Differential expression tests, using discrete pseudo-bulk analysis with Dreamlet (comparing >40 vs. <40 year old donors), and continuous single-cell analysis with kNNage. Analysis is done separately for every MMoCHi cell subset.

**Supplementary Table 11: Gene scores in scHPF factors used for studying variation between age groups.**
Pertaining to Fig. 5C,E,F,H. Table indicating the genes score for each cell type.

**Extended Report 1: Per-cell analysis of MrVI revealed age-related changes in specific cell types and tissues.**
Pertaining to Fig 6. The reports include the age-dependent changes depicted by MrVI, examining each immune lineage and tissue. We provide one report per tissue. **Top row**: UMAP of MrVI embedding. Cells are colored according to their MMoCHi-annotated cell type (right) and by their partition into modules by MrVI (left). The partition into modules is performed by bi-clustering the cell x gene matrix of age-driven LFC estimated by MrVI (giving rise to both gene clusters and to cell clusters). LFC estimates are beforehand smoothed between nearest neighbours in the embedding space (multiplied by normalized connectivities for 20 nearest neighbours). LFC estimates are additionally normalized to an L2 norm of 1 for each gene and each gene is multiplied by the sign of the sum across all cells (genes downregulated and upregulated are treated as the same during biclustering). **Second row**: UMAP with cells coloured by the scores of the respective gene modules. Scores are computed for each cell by summing the inferred LFCs of signature genes that are upregulated in old individuals minus the sum of LFCs of genes downregulated in old individuals. **Third row**: validating module scores in raw expression values. Scores are computed by summing all count normalized and log1p transformed values for upregulated genes with age minus all values downregulated with age. The violin plots show the distribution of module scores for

every MMoCHi-annotated type, separately for subjects >40 and <40 years old. **Fourth row**: (left) UMAP with cells colored by MrVI provided differential abundance analysis estimates. Numbers reported are the log-likelihood difference between all samples from individuals above and below 40 years of age. (right) pseudobulk differential expression analysis, comparing >40 vs. <40 years old within each cell module. **Fifth row**: top genes inside each MrVI gene module. For selection of top genes the normalized LFC values (described above) for all cells inside a cell module is computed and the top 20 genes are selected. Plotted is the average LFC across all cells in a respective cell module.
